# Glomerular Endothelial Cell-Derived Extracellular Vesicles Cross the Basement Membrane to Regulate Podocyte Function

**DOI:** 10.64898/2026.01.28.697274

**Authors:** Janina Kern, Sindhu Thiagarajan, Nina Sopel, Alexandra Ohs, Patricia Luckner, Astrid Bruckmann, Jan Van Deun, Mustafa Kocademir, George Sarau, Silke Christiansen, Christoph Daniel, Mario Schiffer, Stefan Uderhardt, Janina Müller-Deile

## Abstract

**Background:** Small extracellular vesicles (EVs) are nanosized, endosome-derived particles which transfer RNA, proteins, and bioactive molecules to mediate intercellular communication. While EV signaling has been observed in many organ systems, it remains unclear whether glomerular endothelial cell (GEC)-derived small EVs directly interact with podocytes *in vivo* or how they traverse the glomerular basement membrane (GBM).

**Methods:** GEC-derived small EVs were characterized by nanoparticle tracking analysis, electron microscopy, RAMAN spectroscopy and flow cytometry. Cargo composition was analyzed by proteomics, and microRNA (miR) profiling. Functional and structural features were examined using protease, collagenase, adhesion, and multimodal imaging assays. GEC-derived small EV uptake and downstream transcriptional effects were studied in cultured podocytes, while *in vivo* trafficking was assessed by injection of labeled small EVs into transgenic zebrafish larvae under baseline conditions, puromycin-induced damage, and cd2ap-knockdown.

**Results:** GECs released bona fide exosome-like small EVs carrying a highly cell type-specific miR cargo. Small EV transfer to podocytes induced a defined transcriptional response consistent with miR-mediated repression of target genes involved in extracellular matrix organization, cell cycle regulation, and cellular stress responses. Proteomic analyses revealed enrichment of surface proteases and integrin-associated proteins that conferred sustained proteolytic activity and enabled GEC-derived small EV migration through extracellular matrix surrogates. *In vivo*, circulating small EVs traversed the GBM and localized selectively to podocytes in healthy glomeruli, whereas glomerular injury permitted small EV entry into the tubular compartment.

**Conclusion:** These findings provide first *in vivo* evidence that GEC-derived small EVs can cross the GBM and impact on podocytes. By identifying integrin- and protease-dependent mechanisms which facilitate vesicle passage, this study redefines the GBM as a dynamic interface of heterocellular, vesicle-mediated communication.

## Introduction

Extracellular vesicles (EVs) represent a heterogeneous class of membrane-bound particles enclosed by a phospholipid bilayer. They are broadly categorized based on their biogenesis into two major types: small EVs (commonly referred to as exosomes) and ectosomes (also known as microvesicles) (1). Small EVs originate from the endosomal system, forming within multivesicular bodies (MVBs) and are released into the extracellular space via fusion of MVBs or amphisomes with the plasma membrane (2–6). Small EVs carry a selective and dynamic cargo of cytosolic and membrane-associated proteins, RNAs, and lipids which reflect the physiological state and identity of the parent cell.

While much is known about the systemic biodistribution of EVs, their ability to traverse biological barriers remains incompletely understood. In the kidney, EVs are released by diverse resident cell types, including epithelial cells, podocytes, mesangial cells, and endothelial cells and have been detected in urine, implicating them in intrarenal and intercellular communication (7, 8). Increasing evidence supports a role for EVs in the pathogenesis of kidney diseases, where they may propagate injury through delivery of inflammatory cytokines, pro-fibrotic factors, and proteolytic enzymes (9, 10).

The glomerular filtration barrier (GFB) built of fenestrated glomerular endothelial cells (GECs), the glomerular basement membrane (GBM), and podocyte slit diaphragm functions as a size-and charge-selective interface that typically excludes macromolecules and subcellular particles from passing into the urinary space (11). Given their size (∼50–150 nm), small EVs were long thought to be unable to cross this barrier. However, emerging evidence suggests that under certain physiological or pathological conditions, vesicle populations may circumvent or modify this restriction, challenging conventional paradigms of GFB impermeability (12). Importantly, while GEC-to-podocyte communication via small EVs has been suggested in *in vitro* models (13–15), *in vivo* evidence has been lacking.

In this study, we test the hypothesis that GEC-derived small EVs can cross the GBM and modulate podocyte function. We propose that this translocation is facilitated by surface integrin-mediated matrix binding and protease-dependent remodeling of the extracellular matrix. We comprehensively profiled microRNA (miR), and protein cargo of human GEC-derived small EVs and evaluated their impact on podocyte transcriptomics. Integrative analyses allowed us to correlate podocyte gene expression changes with specific components of the small EV cargo. Functional assays, including protease activity measurements and integrin adhesion assays, further delineated the mechanisms of small EV interaction with the GBM. To validate these findings *in vivo*, we employed a transgenic zebrafish model that allows real-time tracking of small EV biodistribution under both physiological and glomerular injury conditions. Our findings suggest a previously unrecognized route of glomerular cell-cell communication and establish a mechanistic framework for small EV-mediated signaling across the GBM. This work provides new insights into the biology of small EVs in renal physiology and its signaling in intraglomerular communication.

## Methods

### Cell culture of GECs and small EV isolation by differential ultracentrifugation

Human GEC (Clonetech, Mountain View, CA, USA) were cultivated in endothelial cell basal media MV2 (PromoCell GmbG, Heidelberg Germany). This medium was supplemented with 5 % FCS, 0.004 ml/ml endothelial cell growth supplement, 10 ng/m L epidermal growth factor, 90 µg/mL heparin, 1 µg/mL hydrocortisone (supplement mix, PromoCell GmbH). 1 % penicillin/streptomycin (Thermo-Fisher Scientific, Waltham, Massachusetts, USA) was added to the medium and cells were maintained at 37°C and 5% CO₂ in a humidified incubator. Flasks, plates and dishes were treated with attachment factor (Cell Systems, Kirkland, Washington, USA) before seeding. At 80% confluency, GECs were starved for 4 h in medium containing 1 % vesicle-depleted FCS (obtained by ultracentrifugation at 110,000 x g overnight) and then treated with media containing 5 % versicle-depleted FCS. After 24h or 48 h cell culture supernatant was harvested and small EVs were isolated by a standard differential ultracentrifugation protocol as follows: The collected supernatant was first centrifuged at 300 × g for 5 minutes at 4°C to remove cells, followed by 2,000 × g for 20 minutes at 4°C to remove debris. The resulting supernatant was then centrifuged at 10,000 × g for 30 minutes at 4°C to eliminate larger vesicles and apoptotic bodies. Before the final ultracentrifugation step which was performed at 150,000 × g for 120 minutes at 4°C using a Beckman Coulter Optima ultracentrifuge with a Type 70 Ti rotor, the supernatant was filtered by a 0.22 µm pore size PES syringe filter (Carl ROTH, Karlsruhe, Germany). The EV-containing pellet was resuspended in PBS for downstream analyses.

### Cell culture of human podocytes

Immortalized human podocytes (gift from Moin Saleem, Bristol, UK) were cultivated in RPMI 1640 media (Gibco, Thermo-Fisher Scientific, Waltham, Massachusetts, USA). The medium was supplemented with 10 % FBS, 1 % penicillin/streptomycin (Thermo-Fisher Scientific, Waltham, Massachusetts, USA) and 0.1 % Insulin-Transferrin-Selen (Gibco, Thermo-Fisher Scientific, Waltham, Massachusetts, USA). Cells were maintained at 33 °C and 5 % CO_2_ in a humidified incubator. For EV isolation, cells were seeded at 80 % confluency and differentiated at 37 °C. At day 8 after differentiation, they were starved for 4 h in medium containing 1 % vesicle-depleted FCS and then treated with medium containing 10 % versicle-depleted FCS for 48 h. Small EVs were isolated by standard differential ultracentrifugation as mentioned before. 15,0000 podocytes were treated with 5×10^9^ GEC-derived small EVs (3.3 10^4^ GEC-derived EVs/ podocyte) for 24 h.

### Nanoparticle tracking analysis

Size distribution and particle concentration of the isolated small EVs were analysed using a nanoparticle tracking analysis system (Malvern Instruments, Worcestershire, UK). Samples were diluted in PBS to a final particle concentration suitable for analysis (typically 10⁷–10⁹ particles/mL). Mean particle size, mode size, and particle concentration were recorded and calculated for further experiments.

### Flow cytometry analysis (FACS) following magnetic bead enrichment

For surface marker profiling, small EVs were first pre-enriched using CD63-coated magnetic beads (Dynabeads, Thermo Fisher Scientific) according to the manufacturer’s instructions. Briefly, small EVs were incubated with CD63-conjugated beads overnight at 4°C with gentle rotation. Bead-bound small EVs were then washed with PBS and incubated with fluorochrome-conjugated monoclonal antibodies against CD9, CD63, and CD81 (e.g., anti-CD9-FITC, anti-CD63-PE, anti-CD81-APC; (Biolegend) for 1 hour at 4°C in the dark. Beads were washed to remove unbound antibodies and analysed using a flow cytometer (Cytoflex; Beckman Coulter). Data were analysed using FlowJo software (version 10.10). To assess the absence of cellular contamination in small EV preparations, FACS surface staining for Calnexin, an endoplasmic reticulum (ER)–resident protein not expected to be present in EVs, was performed in parallel. Small EVs samples prepared as above, were incubated with a Calnexin-CoraLite® Plus 488 primary antibody (Proteintech) under identical staining conditions used for small EV markers in parallel. Following staining, EV samples were acquired using identical acquisition settings as for EV markers. Absence of Calnexin signal in EV preparations confirmed a lack of ER contamination and served as a negative control validating the specificity of EV marker detection. Data were analysed using FlowJo software (version 10.10) and represented as MFI.

### Electron microscopy

For ultrastructural analysis, isolated small EVs were fixed in 2% paraformaldehyde overnight at 4°C and applied to formvar/carbon-coated copper grids 200 mesh (Electron microscopy sciences). After incubation for 20 minutes, grids were contrasted with 2% Uranyless (Electron microscopy sciences) for 10 minutes followed by washing with ddH2O three times and 5 minutes each. This followed by incubation of the grids with lead citrate (0.5%). The grids were washed with ddH2O and air dried for 10 minutes. The grids were mounted on the grid holder and were examined using either a transmission electron microscope (Leo 910, Zeiss, Oberkochen. Germany) operated at 80 kV or a scanning electron microscope (Sigma, Zeiss) operated at 1,5 kV. Images were captured digitally to assess vesicle morphology and structural integrity.

### Stimulated Raman scattering and fluorescence measurements of small EVs

Stimulated Raman scattering (SRS) and fluorescence measurements were performed on a Leica SP8 confocal microscope equipped with an SRS module (Leica Microsystems, Wetzlar, Germany). The system was driven by a picosecond laser source (APE, Germany) providing synchronized pump and Stokes beams, with the Stokes beam fixed at ∼1031 nm and the pump beam tunable to access the desired Raman shifts. The frequency difference between pump and Stokes beams was controlled via the Leica software to target specific vibrational modes. Measurements were carried out using a 40× water-immersion objective (NA = 1.1) for excitation and a 1.4 NA oil condenser for forward-detected SRS signal collection. The pump beam was amplitude-modulated, and the SRS signal was detected in the forward direction using a photodiode detector coupled to a lock-in amplifier (Zurich Instruments, Switzerland). SRS hyperspectral data were acquired in two spectral regions: the fingerprint region (approximately 980–1800 cm⁻¹) and the CH-stretch region (approximately 2800–3200 cm⁻¹). Images were recorded at 512 × 512 pixels, with a pixel dwell time of 15.22 µs, scan speed of 100 Hz, and 2× line averaging. Hyperspectral stacks were used to extract average spectra from selected regions of interest. All spectra were background-corrected and normalized prior to analysis.

Fluorescence imaging was performed on the same system prior to SRS acquisition to localize labeled small EVs. Small EVs were stained with PKH26, a lipophilic membrane dye (excitation ∼551 nm, emission ∼567 nm), and fluorescence images were acquired using the standard Leica fluorescence detection pathway. Fluorescence and SRS measurements were spatially correlated by imaging the same field of view.

### FRET-based and casein-based protease activity assay

Proteolytic activity associated with GEC-derived small EVs was analyzed using a fluorescence resonance energy transfer (FRET)-based assay. Small EVs were thawed on ice and diluted 1:10 in assay buffer (PBS supplemented with Ca²⁺/Mg²⁺; cathepsin substrates were prepared in MilliQ water). A total of 25 μL of diluted small EV sample or buffer control was added to black 384-well plates with black bottoms (Greiner). Fluorogenic FRET substrates (Sensolyte, AnaSpec) were thawed at room temperature, vortexed briefly, centrifuged (15,000 g, 1 min), and diluted 1:100 in assay buffer. Subsequently, 25 μL of substrate solution was added to each well. After gentle mixing, fluorescence signals were recorded at 37 °C for 1 h on a Spectramax M3 plate reader (Molecular Devices) using the “Protease assay.spr” protocol. Background fluorescence (blank wells without EVs) was subtracted from all readings. Quantitative determination of proteases on the intact small EVs were measured using the protease activity assay kit (Abcam) according to the manufacturer’s instructions. The assay uses fluorescence isothiocyanate (FITC)-labelled casein as a substrate. The small EV samples are incubated with the substrate and the fluorescence of the FITC-labeled peptide fragments were measured using a plate reader at the Excitation/Emission = 485nm/530nm. Protease activity was quantified as the increase in fluorescence relative to blank controls.

### Heat inactivation of small EVs

Heat-treated small EVs were used as functional negative controls. Small EV sample aliquots were heat-inactivated prior to the collagenase assay. Aliquots of purified were incubated at 90 °C for 10 min on a heat Block with closed lids to prevent evaporation. Samples were then immediately cooled on ice for 5 min and centrifuged briefly (10,000 × g, 5 min) to remove any precipitated liquid.

### Collagenase activity assay

Collagenase activity of GEC-derived small EVs was quantified using a fluorogenic collagen degradation assay (EnzChek Gelatinase/Collagenase-Assay-Kit, ThermoFischer, Waltham, Massachusetts, USA) according to the manufacturer’s instructions. Briefly, EV preparations were incubated with a quenched fluorescein-labeled collagen substrate, which emits negligible baseline fluorescence when intact. Upon proteolytic cleavage by collagen-degrading enzymes contained within or associated with the EVs, the fluorescent dye is released, resulting in a proportional increase in fluorescence intensity. EV samples were incubated with the substrate in assay buffer at 37 °C, and fluorescence was recorded kinetically at defined intervals using a microplate reader (excitation/emission: ∼490/520 nm). To confirm specificity, parallel reactions containing a broad-spectrum MMP/collagenase inhibitor served as negative controls, while recombinant collagenase was included as a positive control. Fluorescence values were background-corrected, normalized to small EV protein content, and expressed as relative collagenase activity. All measurements were performed in triplicates.

### Adhesion assay for small EV-coated microspheres

Approximately 10^10^ small EVs were passively adsorbed onto 10 ml of Fluoresbrite plain microspheres (2.5% solids-latex, 3 μm YG, Funakoshi, Japan; catalog #17155. Wells of a 96-well V-bottom plate were coated with MAdCAM-1-Fc. Equal volumes of small EVs-coated microspheres were then added to the wells in the presence of either 1 mM MgCl₂/CaCl₂ or 2 mM EDTA. Conditions containing EDTA served to determine background adhesion. After a 20 min incubation at room temperature, plates were centrifuged at 3000 rpm for 5 min. Non-adherent microspheres (accumulating at the bottom of the V-shaped wells) were quantified using an ARVO X-2 Multilabel Reader (PerkinElmer, Japan). The percentage of bound microspheres was calculated as previously described (16).

### RNA isolation, cDNA synthesis, and quantitative PCR

Total RNA from whole-cell lysates was extracted using the innuPREP RNA mini-Kit (iST Innuscreen GmbH, Berlin, Germany) according to the manufacturer’s instructions. For cDNA synthesis, 500 ng of RNA was reverse transcribed with M-MLV Reverse Transcriptase (50,000 U, Promega) in the presence of 5× RT buffer, dNTP mix (Promega), random hexamer primers (Thermo Fisher Scientific), and RiboLock RNase inhibitor (Thermo Fisher Scientific). Reverse transcription was performed at 25 °C for 5 min, 40 °C for 60 min, and 70 °C for 10 min. Quantitative PCR was carried out using the Maxima SYBR Green/ROX qPCR Master Mix (Thermo Fisher Scientific) under the following conditions: 95 °C for 10 min, followed by 40 cycles of 95 °C for 15 s and 60 °C for 1 min, and a final dissociation step (95 °C for 15 s, 60 °C for 1 min, 95 °C for 15 s). All reactions were run in triplicates. Relative expression levels were calculated using the ΔΔCt method. Primer sequences are provided in **supplementary Table S1**.

### Next-generation sequencing of small EV-derived miRs

Next-generation sequencing data was analyzed using the miND® analysis pipeline (17). Overall quality of the NGS data was evaluated automatically and manually with fastQC v0.12 (18) and multiQC v1.14 (19). Reads from all passing samples were adapter trimmed and quality filtered using cutadapt v3.3 (20) and filtered for a minimum length of 17nt. The multiQC report was provided in the supplements (supplementary data).

Mapping steps were performed with bowtie v1.3.0 (21) and miRDeep2 v2.0.1.2 (22), whereas reads were mapped first against the genomic reference GRCh38.p12 provided by (23) allowing for two mismatches and subsequently miRBase v22.1 (24) filtered for miRs of hsa only, allowing for one mismatch. For a general RNA composition overview, non-miR mapped reads were mapped against RNAcentral v23.0 (25) and then assigned to various RNA species of interest. Statistical analysis of preprocessed NGS data was done with R v4.3 and the packages heatmap vNA, pcaMethods v1 and genefilter v1. Differential expression analysis with edgeR v4.0 (26) used the quasi-likelihood negative binomial generalized log-linear model functions provided by the package. The independent filtering method of DESeq2 (27) was adapted for use with edgeR to remove low abundant miRs and thus optimize the false discovery rate (FDR) correction. Additional NGS QC and absolute quantification of miRs was done using miND® spike-ins (28). In order to calculate concentrations of miRs, a linear regression model (y ∼ 0 + x) was used. Quality control evaluation was performed based on the miND® spike-ins as described previously (28). For a sample to pass the calibration quality control checks, the following must be true: 2 or more miND® spike-in core sequences detected, linear model parameters calculated and Pearson correlation coefficient (r-squared) above 0.95. After this miR reads were converted to absolute molecules/uL based on NGS spike-in calibrators. The miND® spike-in consists of seven oligonucleotides that are provided in a specific ratio to cover the broad concentration range of endogenous small RNAs. A unique design of the miND® spike-in reduces sequencing bias and ensures precise quantitation of small. The miND® spike-in sequences are detected in the NGS data along with the endogenous small RNAs. Read counts of the miND® spike-in and endogenous miRs are used to calculate absolute concentrations (molecules/µL).

### Next-generation sequencing (NGS) of podocyte mRNA

Overall quality of the next-generation sequencing data was evaluated automatically and manually with fastQC v0.12 (18) and multiQC v1.14 (19). Reads from all passing samples were adapter trimmed and quality filtered using bbduk from the bbmap package v38.69 and filtered for a minimum length of 17nt and phred quality of 30. Alignment steps were performed with STAR v2.7 (29) using samtools v1.9 (30) for indexing, whereas reads were mapped against the genomic reference GRCh38.p13 provided by Ensembl (23). Assignment of features to the mapped reads was done with htseq-count v0.13 (27). Differential expression analysis with edgeR v3.40 (26) used the quasi-likelihood negative binomial generalized log-linear model functions provided by the package. The independent filtering method of DESeq2 (27) was adapted for use with edgeR to remove low abundant genes and thus optimize the FDR correction.

### GO enrichment analysis

Differentially expressed mRNAs based on a FDR cutoff <= 0.05 were used in a GO-term enrichment analysis. Enriched biological processes were identified by using the Kolmogorov Smirnov (KS) test with the unadjusted p-values from the differential expression analysis as rank information with the tool topGO (31).

### Heatmaps from miR NGS of small EVs

Data was based on RPM normalized reads and scaled using the unit variance method for visualization in heatmaps. Clustering was done using the average method of heatmap calculating the distances as correlations. An additional filter was introduced to increase the robustness: only miRs that showed an RPM in at least 1/ n (groups) percent of samples (e.g. with 4 groups, the miR had to have an RPM value above 5 in at least 25% of the samples). This removed miRs that had a high CV but are only expressed in a too small number of samples to bear any statistical significance or biological relevance.

### miR-mRNA interaction analysis

Two different approaches were chosen to perform this analysis: We connected the top 20% abundant miRs of the GEC-derived small EVs with the 40 mRNAs that are significantly regulated from the corresponding contrast. We first performed target prediction of the Top 37 miRs resulting in ca. 4,400 predicted targets. Of these, 20 were part of the set of significantly regulated mRNA comparing treated versus untreated podocyte samples. As an alternative angle, we used the miRs which were significantly enriched in GEC-derived small EV versus podocyte-derived small EVs (59 miRs) and performed miR-mRNA interaction analysis. The tool miRtap (32) allowed to combine the results of 5 different miR-mRNA interaction databases (targets are aggregated from 5 most commonly cited prediction algorithms: DIANA (33), Miranda (34), PicTar (35), TargetScan (36), and miRDB (37). By providing the miRBase ID of individual miRs, a predictions table containing the ranks for each target gene as well as the aggregated rank product were reported. The number of target genes were filtered by specifying a minimum number of 3 databases where the target gene must have been be found.

### Small EV sample preparation and LC-MS/MS

For proteomic analysis, 3.17×10^8^ small EVs were lyophilized and sample preparation was performed using EasyPep™ MS Sample Prep Kit (Catalog Number A40006, Thermo Fisher Scientific) according to the manufacturer’s instructions with minor changes. Briefly, sEVs were lysed, reduced, alkylated and subjected to digest with 1 µg Trypsin/LysC-mix over night at 37°C. After peptide clean-up, samples were vacuum dried and reconstituted in 0.1 % formic acid. Separation of peptides (0.2 µg) by reversed-phase chromatography was carried out by a proteoElute system which was equipped with a C18 ProteoTrap preconcentration column (250 μm i.D. x 5 mm, 3 µm particle size, Bruker Daltonics) in front of a PepSep Ultra C18 nano column (75 μm i.D. x 250 mm, 1.5 µm particle size, Bruker Daltonics). A linear gradient of 4 % to 40 % acetonitrile in 0.1 % formic acid over 22 min was used to separate peptides at a flow rate of 250 nl/min. The LC-system was coupled on-line to a timsTOF HT System (Bruker Daltonics) via a CaptiveSpray 2 nanoflow electrospray source (Bruker Daltonics). Data were acquired under dia-PASEF mode with a MS1 scan range of 100–1700 m/z, MS2 scan range of 400–1000 m/z and an ion mobility range from 0.65 to 1.45 1/K0 (inverse reduced ion mobility) via the Compass HyStar 6.4 acquisition and processing software (Bruker Daltonics).

Raw data were processed using Bruker ProteoScape (BSP version 2026) software with an implemented Spectronaut software (SN v20.2) and searched against the human swissprot database (download October 2025). Search parameters were set up as follows: directDIA+ workflow, enzyme specificity trypsin, a maximum of two missed cleavage allowed, deamidation (N, Q), acetylation (Prot N-term) and oxidation (M) as variable modifications and carbamidomethylation (C) as fixed modification. Precursor and protein (global) Q-value Cutoff were set to 0.01. Venn diagrams were drawn using the following online tool: https://bioinformatics.psb.ugent.be/webtools/Venn/

### Zebrafish embryo handling

Tg(wt1b:eGFP) zebrafish were set up to breed at a ratio of 3 to 2 females to males overnight under standard conditions at 28 °C. Eggs were collected the next morning in embryo raising medium (ERM) as previously described (38) and transferred to a 28 °C incubator. Unfertilized and abnormally formed embryos were removed in the following 48 h. 24 h post fertilization (hpf) the embryos were treated with N-Phenylthiourea (Sigma-Aldrich, Burlington, Massachusetts, USA) to prevent pigmentation to limit background fluorescence to the minimum. Larvae were screened for a positive Wt1b_GFP signal and larvae with no signal were sorted out.

Cd2ap morpholino (MO) was injected into one to two cell stage of zebrafish eggs as published before (39). In brief cd2ap-MO, which was diluted in injection buffer (1:1), was injected at final concentrations of 50 μM in a total volume of 4.6 nL. MO sequences for *cd2ap* were designed and ordered from GeneTools (Philomath, OR) as follows: 5′-CATACTCCACCACCACCTCAACCAT-3′. Sequence for the control MO was 5′-CCTCTTACCTCAGTTACAATTTATA-3′. The cd2ap-MO sequence was blasted to ensure that no off-target annealing occurred for sequence matches >14 nt, and we excluded cross hybridization with the cRNA constructs. Puromycin (PAN)-treatment of zebrafish larvae to induce glomerular injury was performed as published before (40). In brief, at 46 hpf zebrafish were treated with either 6 mg/ml PAN by adding the drug into the fish medium with 1% DMSO. Fish were kept in this PAN-containing medium until 96 hpf. DMSO, which was added to the fish water in the same concentration, served as control

### Cardinal vein injection of small EVs in zebrafish larvae

Cardinal vein injections were conducted 72 h post fertilization with the microinjector Nanoject II (Drummond Scientific Company, Broomall, Pennsylvania, USA) as previously described (15, 41). Capillary needles for injection were cut back to obtain an opening of about 20 µm. Human-derived fluorescently labeled small EVs were stained with PKH26 red fluorescent cell linker kit (Sigma-Aldrich, Burlington, Massachusetts, USA) for EV membrane labelling. The labeled small EVs were filled into a needle containing mineral oil. The larvae were immobilized by treating them with 0.4 % Tricain (Sigma-Aldrich, Burlington, Massachusetts, USA) and placed in an 1.5 % agarose mold. 4,6 nL of sample were injected into each embryo into the cardinal vein which approximately matches 200,000 small EVs per larvae. After injection, the embryos were transferred back to PTU-containing ERM and kept on 28 °C.

### Pronephroi isolation

The approach for the isolation of the pronephroi was published before (42). Zebrafish larvae were incubated in 0.2 % Tricain (Sigma-Aldrich, Burlington, Massachusetts, USA) with 10 mM DTT for 1 h at room temperature at 96 hpf for degradation and to enable digestion. Afterwards they were washed with PBS-Ca^2+^-Mg^2+^ (PBS+) three times while removing as much PBS+ as possible after the last washing step. The larvae were digested by incubating them with 100 µl collagenase (Sigma-Aldrich, Burlington, Massachusetts, USA) diluted in PBS+ to a concentration of 5 mg/mL for 2 h at 30 °C. The reaction was stopped with 1 mL of DMEM supplemented with 10 % FCS. The mixture was carefully pipetted up and down, transferred to a 3 cm dish and more DMEM with FCSwas added. The pronephroi were identified by morphology on a stereomicroscope (Leica S9D) and collected with a 20 µl pipet.

### Cryosections of zebrafish larvae

Zebrafish larvae were sacrificed by rapid cooling method. Fixation was performed by using 2 % PFA for 2 h at room temperature. The embryos were washed 3 times with PBS for 5 minutes at room temperature. For dehydration, 15 % sucrose diluted in PBS was added for 1 h at room temperature or 4°C overnight. Afterwards, zebrafish larvae were embedded in a 1:2 mixture of 30 % sucrose with tissue tec (Sakura Finetek, Tokyo, Japan) (15 % sucrose end-concentration) and frozen on dry ice or liquid nitrogen. Cryo slices were done with 10 µm thickness.

### Fluorescent imaging

Embryos were immobilized with tricaine and transferred to 96 well plates with a glass bottom. Zebrafish were screened at 96 h post fertilization in a fluorescence microscope from Acquifer IM (Acquifer Imaging GmbH, Heidelberg, Germany) for identification of successfully injected larvae.

### Statistics

Multiple testing adjustment to reduce the false discovery rate (FDR) was done using p-value adjustment methods like Benjamini Hochberg with an arbitrary cutoff for low expressed miRs prior analysis. In order to give the highest sensitivity to our analysis, we implemented a method of removing low read count miRs from the data set until a statistically relevant set of significant results remained (independent filtering). All data are shown as mean ± SEM and were compared by ANOVA or T-test to analyze for statistical significance. Experiments were performed at least three times. If *p* < 0.05, experimental findings were considered significant. All photomicrographs were made at similar intensities and backgrounds.

## Results

### GECs secrete small EVs with characteristics of exosomes

GEC-derived small EVs isolated using differential ultracentrifugation and pore size filtration exhibited characteristics consistent with small EVs, as described before (43). Nanoparticle tracking analysis revealed a median size distribution between 100 and 150 nm, corresponding to the small EV category according to the MISEV 2023 classification guidelines (**Fig. 1A**). Scanning electron microscopy showed vesicles with the typical intact morphology characteristic of small EVs (**Fig. 1B**). To verify the molecular composition of GEC-derived small EVs, we performed correlative fluorescence Stimulated Raman Scattering (SRS) imaging on red fluorescently labeled EV preparations. SRS microscopy is a label-free vibrational imaging technique that amplifies weak Raman scattering signals by driving specific molecular bonds into resonance using two synchronized laser beams (pump and Stokes). When the energy difference between the lasers matches a molecular vibrational frequency, energy transfer occurs, producing a strong, background-free signal which is linear with molecular concentration. Fluorescence–SRS correlative imaging showed complete spatial overlap between the small EV-derived red fluorescence and the SRS signal (**Fig. 1C**). SRS hyperspectral measurements of isolated small EVs revealed vibrational signatures consistent with lipid- and protein-rich small EVs. In the fingerprint region (980–1800 cm⁻¹), the spectra exhibit prominent bands near 1650–1660 cm⁻¹, corresponding to the amide I vibration of proteins, as well as a band around ∼1445 cm⁻¹ attributed to CH₂ bending modes of phospholipid acyl chains. Additional features observed in the 1130–1160 cm⁻¹ range are assigned to C–C and C–N stretching modes associated with lipid backbones and protein side chains (**Fig. 1D**). In the CH-stretch region (2800–3200 cm⁻¹), the spectra display a lipid-dominated profile, with strong CH₂ symmetric and asymmetric stretching bands between ∼2845–2890 cm⁻¹ and a CH₃ stretching band around ∼2930–2950 cm⁻¹ (**Fig. 1D**). For further characterization of GEC-derived small EVs their surface markers were analyzed by bead-based flow cytometry, using capture beads coated with antibodies against small EV surface proteins to immobilize the vesicles, followed by detection with fluorescently labelled antibodies targeting tetraspanins. This approach enabled the characterization of GEC-derived small EVs by FACS, confirming the presence of classical exosome markers CD63, CD9, and CD81 on their surface (**Fig. 1E; lower panel**). Small EV depleted FCS served as a control (**Fig. 1E; upper panel**). The absence of calnexin, an endoplasmic reticulum-resident chaperone, confirmed minimal ER contamination of purified GEC-derived small EV preparations (**Fig. 1E**). Thus, GECs are capable to generate and secrete small EVs that display key features of exosomes described in various biological systems.

**Figure 1:**
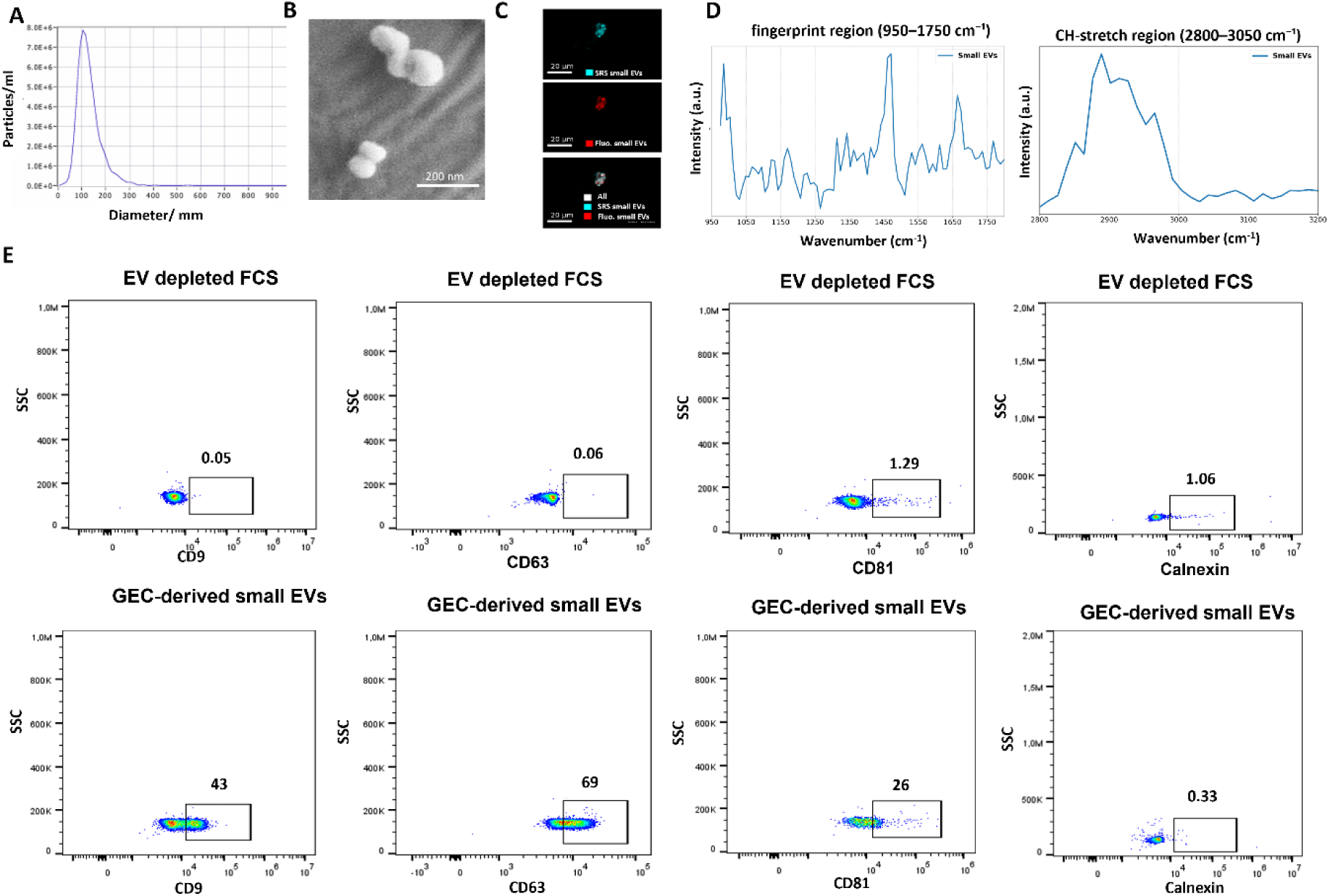
Basic characterization of small EVs isolated form GEC cell culture supernatant. A: Nanoparticle tracking analysis (NTA) analysis of GEC-derived small EVs reveals that the EVs have a median size of 100 nm. B: Scanning electron microscopy of GEC-derived small EVs showing a median size of 100 nm. C: Representative fluorescence stimulated Raman scattering (SRS) overlay image of GEC-derived small EV clusters. Small EVs labeled with PKH26 are shown in red (fluorescence, excitation 551 nm), while the SRS signal acquired at 2940 cm⁻¹ (CH₂ stretching vibration) is shown in cyan, highlighting lipid-rich vesicle structures. Scale bar: 20 µm. D: Corresponding SRS spectra extracted of GEC-derived small EVs, showing the fingerprint region (980-1800 cm⁻¹) and the CH-stretch region (2800–3200 cm⁻¹). Spectra reveal a prominent peak at ∼1650 cm⁻¹ in the fingerprint region, consistent with amide I vibrations of vesicle-associated proteins and/or C=C stretching of unsaturated lipids, and strong CH₂ symmetric and asymmetric stretching signals in the 2850–2950 cm⁻¹ region, confirming a lipid-rich EV membrane composition. E: FACS analysis with anti-CD9 detection antibody (first column), anti-CD63 detection antibody (secondary column), anti-CD81 detection antibody (third column) and calnexin (forth column). Small EVs were captured with an anti-CD63 capture antibody on beads prior to FACS analysis. First row shows small EV depleted FCS that was used in cell culture media for GECs, second row shows results from GEC-derived small EVs isolated form cell culture supernatant. In case of anti-CD63 detection antibody EVs were captured with an anti-CD63 capture antibody on the beads. Additional CD63 on the captured EVs gave the detectable signal. Abbreviations: EVs: extracellular vesicles, FACS: fluorescence-activated cell sorting, GEC: glomerular endothelial cells, NTA: nanoparticle tracking analysis Abbreviations: EVs: extracellular vesicles, GEC: glomerular endothelial cells.

### Cell type-specific miR signatures distinguish GEC- from podocyte-derived small EVs

To further characterize the small RNA cargo and to compare it with other glomerular cell–derived small EV cargo, we performed next-generation sequencing of mRNAs isolated from GEC- and podocyte-derived small EVs, enabling absolute quantification through the use of miND® spike-ins (28) (**Fig. 2A**). RNA sequencing revealed that GEC-derived small EVs are dominated by rRNA, but consistently contain diverse classes of regulatory RNAs, including miRNA (miRs), tRNA fragments, and piRNAs, with smaller contributions from mRNAs and lncRNAs, consistent with selective RNA packaging into small EV (**supplementary Fig. 1A, B**). In total, 350–450 unique miRs were detected in GEC-derived small EVs, compared with 600–750 in podocyte-derived small EVs. Heatmap clustering of miRs detectable in small EVs revealed distinct profiles for GEC- and podocyte-derived small EVs, indicating that each cell type packages a specific set of miRs (**supplementary Fig. 2**). By far the most abundant miR in both GEC- and podocyte-derived small EVs was miR-21-5p, which is intrinsically one of the most highly expressed in the kidney (**Fig. 2B**) (44). Among the top 100 enriched species were cell-type-associated miRs previously described in glomerular cells, including miR-378a, miR-192-5p, miR-143-3p, miR-26b, miR-200b, and miR-146 (**supplementary table 1 and 2**). Differential expression analysis identified 130 miRs with significantly different abundances between GEC- and podocyte-derived small EVs (FDR < 0.05) (**Fig. 2C**). For example, miR-143 and miR-26b were enriched in GEC-derived small EVs, whereas miR-200b, miR-146, miR-200a, miR-155, and miR-378a were more abundant in podocyte-derived small EVs. miND® spike-ins were used to determine the absolute concentrations of miRs detected by NGS. Comparison of small EV cargo between GECs and podocytes revealed that GEC-derived small EVs are selectively enriched in specific miRs, including miR-9901, miR-122-5p, miR-39860, miR-224-5p, and miR-214-3p (**Fig. 2D**). Conversely, miRs such as miR-429, miR-138-5p, miR-96-5p, miR-183-5p, and miR-102-5p were significantly depleted in GEC-derived small EVs relative to podocyte-derived EVs (**Fig. 2E**). These findings indicate a cell type-specific sorting of miRs into small EVs with yet unclear biological implication.

**Figure 2:**
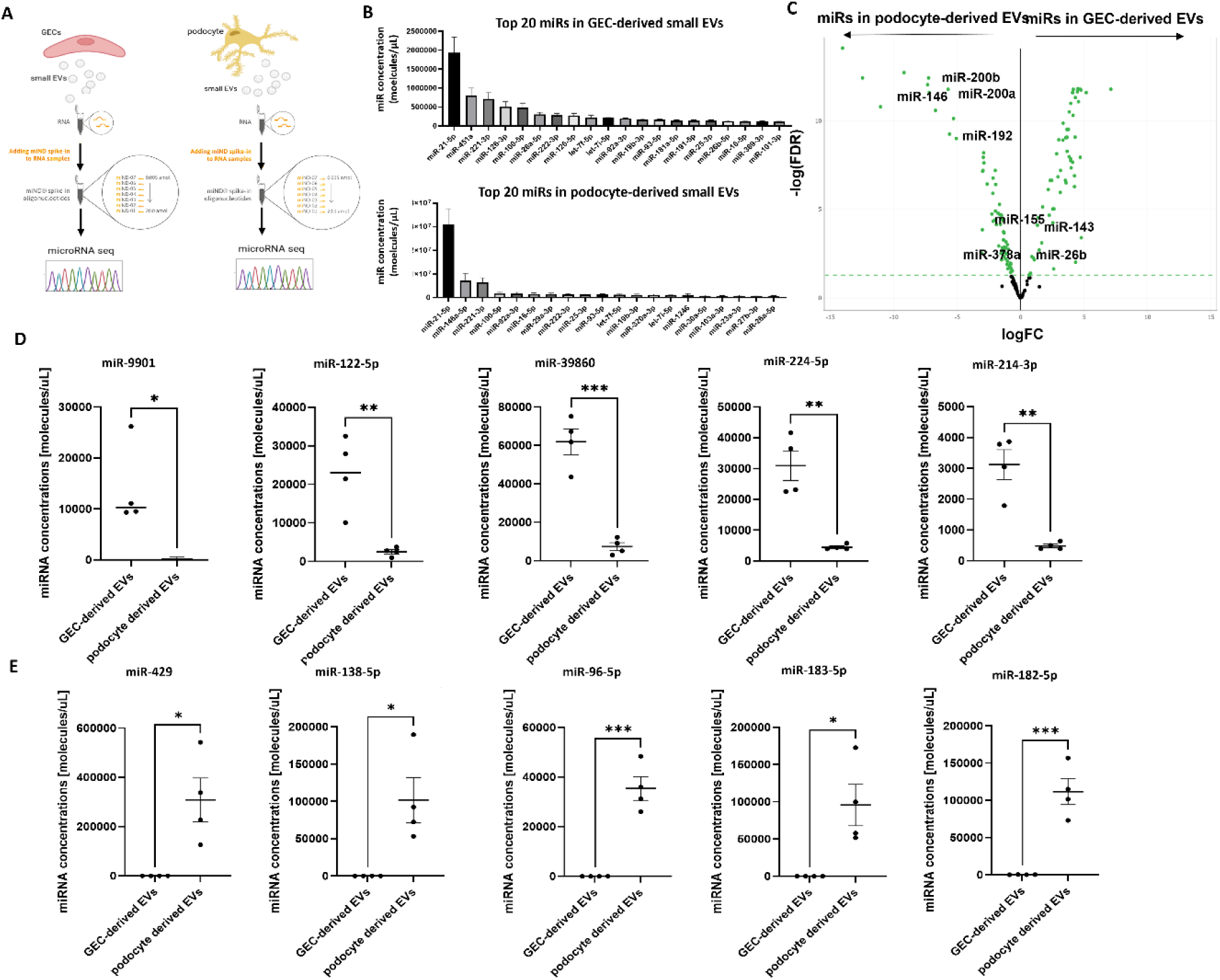
miR profiling of GEC-and podocyte-derived small EVs. A: Schematic illustration of the experimental setup. Small EVs were isolated form GEC and podocyte cell culture supernatant. RNA was isolated from small EVs and samples were spiked with miND before sequencing. miR analysis of GEC-derived and podocyte small EV cargo was performed. B: Most abundant miRs in GEC- and phorocyte-derived small EVs. Abundance of Upper graph: Top 20 miRs in GEC-derived small EVs. Data is given in miR concentration (molecules/µl) as mean +/- SEM after normalization with spike in controls. Lower graph: Top 20 miRs in podocyte derived small EVs. Data is given in miR concentration (molecules/µl) as mean +/-SEM after normalization with spike in controls. C: Volcano blot visualizing the relation of the logFC of miR abundance in GEC-derived small EVs versus miR abundance in podocyte-derived small EVs and the statistical significance of this change. Statistical significance is expressed as false discovery rate (FDR) according to Benjamini and Hochberg. Highlighted are miRs described in glomerular cells before. D: Top 5 upregulated miRs in GEC-derived small EVs compared to podocyte-derived small EVs (FDR < 0.05). miR reads are converted to absolute molecules/uL based on NGS spike-in calibrators. Significance of the difference between miR concentration in GEC-derived and podocyte derived small EVs was calculated with t-test, n = 4, * p<0.05, ** p< 0.001, *** p< 0.001. E: Top 5 downregulated miRs in GEC-derived small EVs compared to podocyte-derived small EVs (FDR < 0.05). miR reads are converted to absolute molecules/uL based on NGS spike-in calibrators. Significance of the difference between miR concentration in GEC-derived and podocyte derived small EVs was calculated with t-test, n= 4, * p<0.05, ** p< 0.001, *** p< 0.001. Abbreviations: EVs: extracellular vesicles, GEC: glomerular endothelial cells, miR: microRNA, NGS: next generation sequencing, SEM: standard error of mean.

### GEC-derived small EVs modulate transcriptional networks in podocytes

To distinguish intracellular CD63-positive compartments from extracellularly released small EVs and to directly visualize EV secretion dynamics in GECs, we employed a pH-sensitive CD63 reporter that emits red fluorescence in acidic lysosomal compartments and green fluorescence in neutral extracellular environments (**Fig. 3A, B**). To compare active vesicle secretion from GECs to passive exposure, we established a transwell co-culture system in which GECs express a green-labelled small EV reporter, allowing us to monitor small EVs released by living cells and their association with podocytes. This setup allows assessment of small EVs secreted *in situ*, as opposed to direct addition of isolated small EVs (**Fig. 3C**). Co-culture of reporter-expressing GECs with podocytes enabled continuous small EV release and transport to podocytes, thereby more closely recapitulating physiologic intercellular communication within the glomerulus (**Fig. 3D, supplementary video 1**).

**Figure 3:**
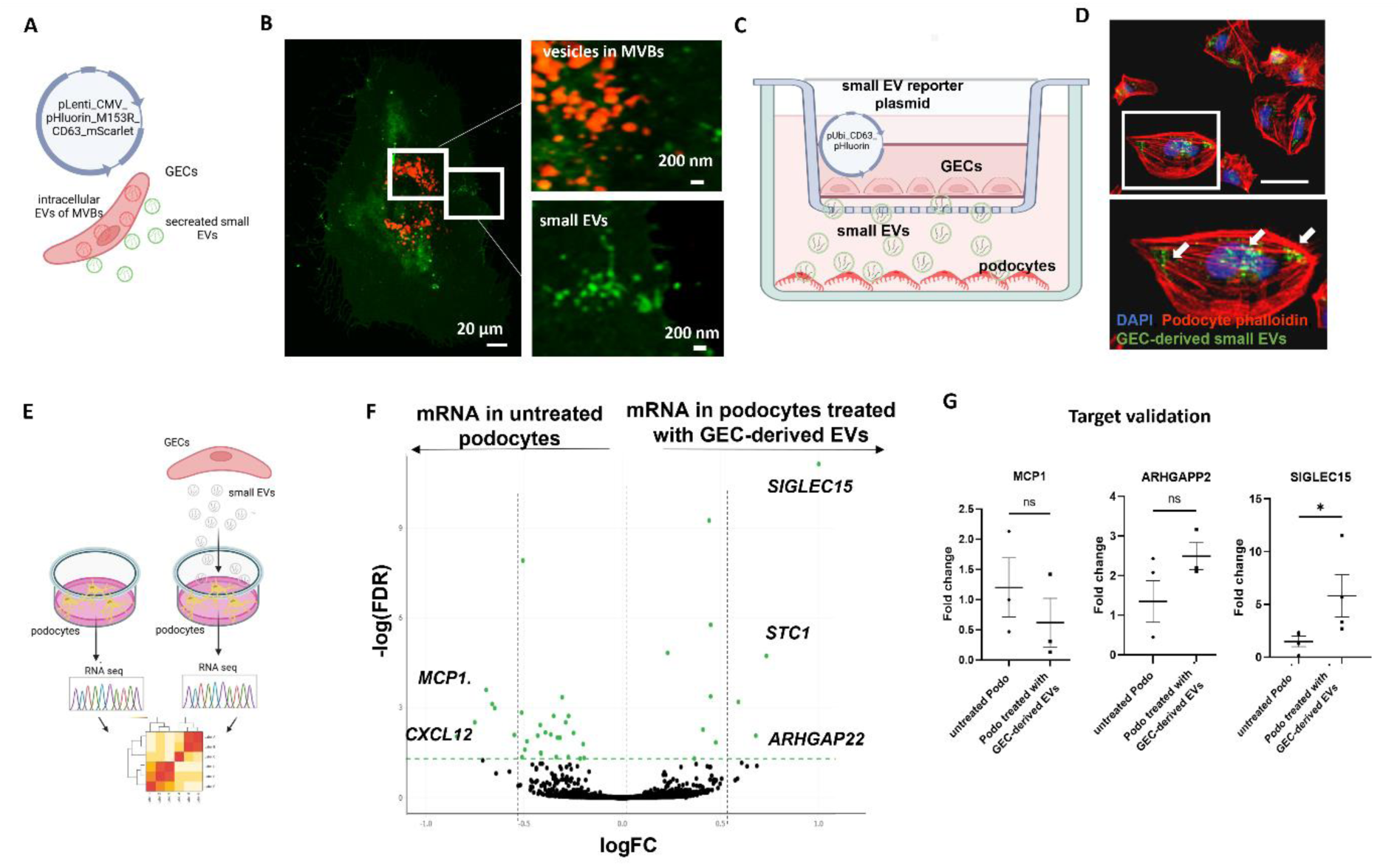
Changes in mRNA expression in podocytes after treatment with GEC-derived small EVs. A: Schematic illustration of GECs electroporated with pLenti_CMV_pHluorin_M153R_CD63_mScarlet plasmid enabling the visualization of intracellular CD63 positive small EVs in acidic pH in red and extracellular in neutral pH in green. B: Immunofluorescent imaging of GECs electroporated with pLenti_CMV_pHluorin_M153R_CD63_mScarlet plasmid showing red CD63 positive small EVs in multivesicle bodies (MVBs) and green CD63 positive small EVs during secretion and outside the cell. C: Schematic illustration of the experimental setup. GECs were electroporated with a small EV reporter plasmid, labeling small EVs in green. Electroporated GECs were then seeded on the upper chamber of a transwell insert, while podocytes were seeded in the lower chamber. D: Immunofluorescent images of podocytes from the transwell co-culture experiment setting (C). After 48 h co-culture of podocytes with the GEC transfected with the small EV-reporter plasmid, podocytes were PFA-fixed and stained with phalloidin to stain the actin structures. The prepared slides were imaged using the confocal microscope (40x). Internalized small EVs are visible within the podocyte cytosol (green fluorescence indicated by white arrows). Scale bar of 40 µm. E: Schematic illustration of the experimental setup. Small EVs were isolated form GEC culture supernatant. Small EVs were then cultured on podocytes for 24 h. NGS-sequencing of mRNA of EV treated and untreated podocytes was performed. F: Vulcano plot visualizing the relation of the logFC of mRNA abundance in podocytes treated with GEC-derived small EVs versus untreated podocytes and the statistical significance of this change. Statistical significance is expressed as false discovery rate (FDR) according to Benjamini and Hochberg to account for multiple comparisons. G: Expression of mRNA for *MCP1. ARHGAPP2* and *SIGLEC15* after exposure of cultured human podocytes to GEC-derived small EVs for 48 h. mRNA expression was normalized to *HPRT.* Differences were calculated between means ± SEM. n = 4, * p < 0.05; n.s. = not significant. Abbreviations: EVs: extracellular vesicles, GEC: glomerular endothelial cells, h: hour, NGS: next generation sequencing.

The transcriptional impact of GEC-derived small EVs on podocytes, was analyzed by exposing cultured human podocytes to GEC-derived small EVs for 24 hours and performing differential gene expression analysis (**Fig. 3E**). EV treatment induced consistent transcriptional changes characterized by both up- and downregulation of defined gene sets, indicating that GEC-derived small EVs simultaneously activate and repress distinct transcriptional programs in podocytes (**Fig. 3F**). In total, 40 genes were significantly differentially expressed in podocytes treated with GEC-derived small EVs compared with untreated controls. Notably, *SIGLEC15* and *ARHGAP22* were upregulated, whereas *MCP*1 was downregulated, suggesting a shift towards a podocyte-protective transcriptional profile (**Fig. 3F, G**). The most strongly upregulated genes in podocytes after treatment with GEC-derived small EVs were *SIGLEC15*, *STC1*, *PREP*, *ARHGAP22* and *EMP1* (**supplementary Fig. 3A**). Top 5 downregulated mRNA in podocytes after treatment with GEC-derived small EVs were *CXCL12*, *MCP1*, *MAP3K7CL*, *TNFRSF9* and *COL11A1* (**supplementary Fig. 3B**). Gene ontology enrichment analysis of the most variable transcripts revealed significant overrepresentation of biological processes related to development, cell proliferation, and regulation of apoptosis (**supplementary Fig. 4**). These findings indicate that GEC-derived small EVs modulate fundamental cellular programs in podocytes, including survival, differentiation, and structural organization.

### Integrative analysis reveals functional GEC-derived miR–podocyte mRNA interactions

To investigate the functional consequences of GEC-derived small EV miR cargo on podocytes, we divided GEC-derived small EV preparations into two groups: One underwent miR profiling, while the other was applied to podocyte cultures (**Fig. 4A**). To predict putative podocyte target genes of GEC-derived small EV miRs, we employed the miRtap tool (32), which integrates five major interaction databases (DIANA, Miranda, PicTar, TargetScan, and miRDB) and reports aggregated target ranks filtered for genes supported by at least three sources. We connected the top 20% most abundant miRs in GEC-derived small EVs with the 40 significantly regulated mRNAs identified in podocytes treated with GEC-derived small EVs compared with untreated controls. In a complementary approach, we also used miRs significantly enriched in GEC-versus podocyte-derived small EVs for interaction analyses.

**Figure 4:**
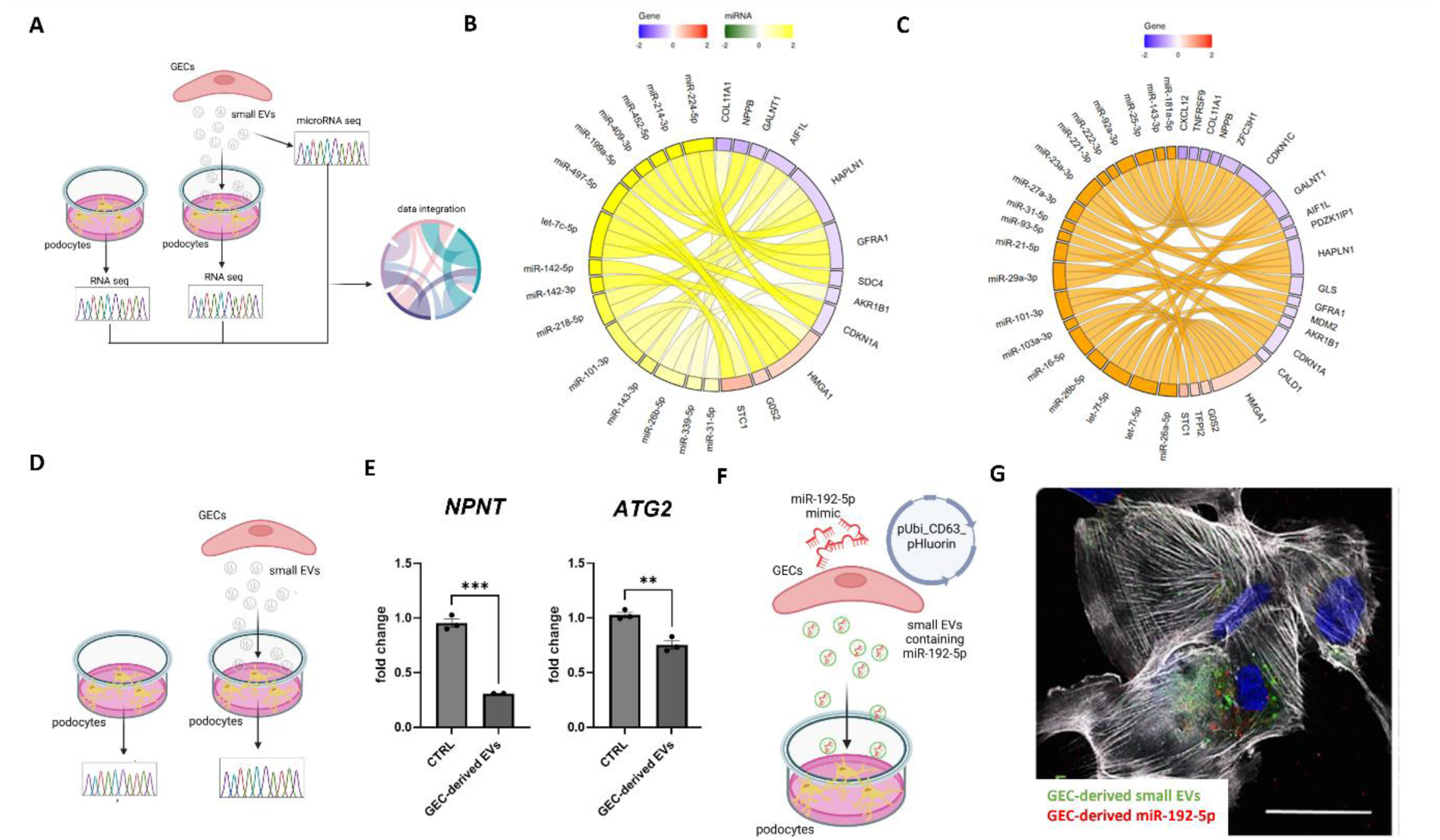
Interaction analysis between miRs of GEC-derived small EVs and podocyte mRNAs. A: Schematic illustration of the experimental setup. Small EVs were isolated from GEC culture supernatant. Small EVs were than cultured on podocytes for 24 h. miR analysis of GEC-derived small EV cargo as well as NGS-sequencing of mRNA of small EV treated and untreated podocytes was performed. Significant enriched miRs were correlated with mRNA targets regulated in podocytes after small EV treatment. B: Chord diagram visualizing predicted miR-mRNA interactions based on your filtered dataset. Left half = miRs (yellow–green scale); right half = predicted target genes (red–blue scale). Each ribbon represents a predicted regulatory interaction between a specific miR and its target gene. miRs data of GEC-derived small EVs was used as log2 fold change (yellow = upregulated, green = downregulated, range −2 to +2 as compared to miRs from podocyte derived small EVs). Podocyte genes were used as log2 fold change (red = upregulated, blue = downregulated after treatment with GEC-derived small EVs). C: Chord diagram visualizing predicted miR-mRNA interactions based on your filtered dataset. Predicted target genes (red-blue scale). Each ribbon represents a predicted regulatory interaction between a specific miR and its target gene. Top 20% abundant miRs of the GEC-derived small EVs were used and podocyte genes were used as log2 fold change (red = upregulated, blue = downregulated after treatment with GEC-derived small EVs). D: Schematic illustration of the experiment of the results form B-E. Small EVs were isolated from GECs and cultured on podocytes for 24 h. Untreated podocytes served as control. RNA was isolated from both groups for qPCR analysis. E: Fold change in podocyte *NPNT* mRNA expression after treatment with GEC-derived small EVs for 24 h. Fold change is given compared to untreated podocytes together with SEM. Expression is normalized to *GAPDH*. *** p < 0.001. n = 3. F: Fold change in podocyte *ATG2A* mRNA expression after treatment with GEC-derived small EVs for 24 h. Fold change is given compared to untreated podocytes. Expression is normalized to *GAPDH*. ** p < 0.01. n = 3. F: Schematic illustration of the experiment shown in panel G. GECs were electroporated with a red fluorescent miR-192 mimic and a small EV reporter plasmid that labels secreted small EVs in green. EVs were collected from the culture supernatant and applied to podocytes. Immunofluorescence imaging was performed to assess the uptake of small EVs and the delivery of miR-192 cargo into podocytes. G: Immunofluorescent image of podocytes labeled with their cytoskeleton (phalloidin, white) and treated with GEC-derived small EVs. GEC were electroporated with a red fluorescent miR-192 mimic and with a green fluorescent exosome reporter plasmid prior to small EV isolation form the supernatant as shown in (F). Both green fluorescent GEC-derived small EVs as well as red fluorescent miR-192 are visible within podocytes. Scale bar 20 µm. Abbreviations: EVs: extracellular vesicles, GEC: glomerular endothelial cells, h: hour, miR: microRNA.

Most of the differentially expressed miRs were upregulated in GEC-derived small EVs, while many of their predicted targets were downregulated in podocytes, consistent with canonical miR-mediated repression. A chord diagram visualizing these interactions highlighted several miRs which were either specifically upregulated in GEC-derived small EVs versus podocyte derived small EVs (**Fig. 4B**) or among the top 20% of miRs highest abundant in GEC-derived small EVs (**Fig. 4C**) as regulatory hubs targeting multiple genes. Conversely, *HMGA1, CDKN1A*, and *SDC4* emerged as convergence nodes for several distinct miRs, suggesting coordinated regulation of pathways related to cell cycle control *(CDKN1A, HMGA1*), extracellular matrix organization (*COL11A1, GALNT1, HAPLN1*), and cellular stress responses (*GOS2, STC1, AIF1L*) (**Fig. 4B, C**). Furthermore, miR-192-5p target genes *NPNT*, a podocyte-derived extracellular matrix protein and *ATG2* were downregulated in podocytes after treatment with GEC-derived small EVs (**supplementary Fig. 5, Fig. 4D, E**). To directly visualize this intercellular transfer of miR containing small EVs, GECs were co-transfected with a CD63-pHluorin construct and a red fluorescent miR-192-5p mimic. Podocytes were then exposed to green fluorescent small EVs carrying red fluorescent miR-192-5p, isolated from GEC culture supernatants (**Fig. 4F**). Immunofluorescence imaging confirmed the uptake of miR-192-containing small EVs by podocytes, providing direct evidence of miR transfer via small EVs (**Fig. 4G**). Thus, we could directly show that the heterocellular transfer of miR from GECs to podocytes via small EVs.

### Protein profiling of GEC-derived small EV cargo

To comprehensively characterize the cargo of GEC-derived small EVs proteomic profiling was performed (**Fig. 5A**). Proteomic analysis of five independent batches of GEC-derived small EVs identified 2,859 proteins in total, with 467 proteins reproducibly detected across all preparations, defining a robust and conserved GEC-EV protein core (**Fig. 5B, supplementary table 3**). In line with the FACS analysis, several exosome markers were detectable on our GEC-derived small EVs (**table 1**). Furthermore, several proteases were present in/on GEC-derived small EVs (**table 2**). Typical non-exosomal contaminants such as histone proteins, cytochrome or ER proteins were not detectable. Pathway enrichment analysis revealed proteins involved in VEGFA-VEGFR signaling, vesicle-mediated transport, regulation of vesicle-mediated transport and extracellular matrix organization were enriched in 5 different batches of GEC-derived small EVs (**Fig. 5B, C**).

**Figure 5:**
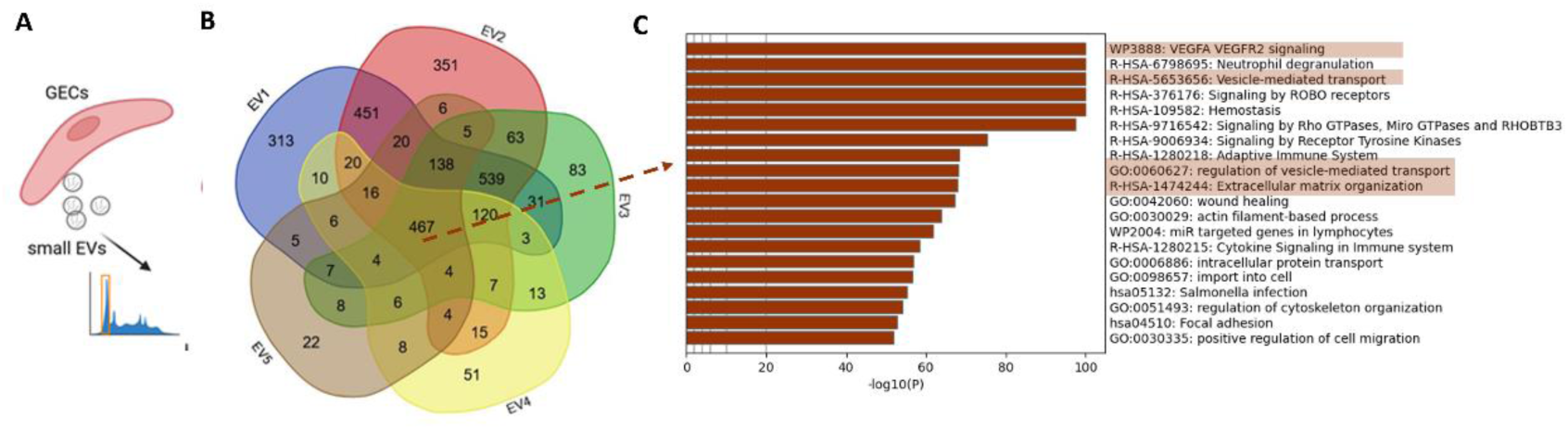
Characterization of GEC-derived small EV RNA cargo and protein cargo. A: Schematic illustration of the experiments to characterize protein cargo isolated from small EVs of cell culture supernatant of GEC. B: Venn diagram illustrating the overlap of proteins identified by mass spectrometry in five independent preparations of GEC-derived small EVs (EV1-EV5). A total of 2,859 proteins were detected across all samples, with 467 proteins consistently identified in all five small EV preparations, defining a conserved core proteome of GEC-derived small EVs. The remaining proteins were variably detected between individual preparations, reflecting biological and/or technical variability. C: Bar plot showing significantly enriched biological pathways and processes identified from the 467 proteins consistently detected across all five independent GEC-derived small EV preparations. Enrichment analysis was performed using pathway and Gene Ontology databases, and significance is expressed as −log10(P value). The conserved small EV protein core was enriched for pathways related to VEGFA-VEGFR2 signaling, vesicle-mediated transport and its regulation, extracellular matrix organization, cytoskeletal dynamics, and cell migration.

**Table 1.** Exosome maker detected via proteomic analysis of GEC-derived small EVs.

|  |
| --- |
| **Tetraspanins**<br>CD63 – CD63 molecule<br>CD9 – CD9 molecule<br>CD81 – CD81 molecule<br>CD82 – CD82 molecule<br>CD151 – CD151 molecule |
| **ESCRT components / exosome biogenesis**<br>TSG101 – Tumor susceptibility gene 101<br>PDCD6IP (ALIX) – Programmed cell death 6 interacting protein<br>CHMP3 – Charged multivesicular body protein 3<br>CHMP6 – Charged multivesicular body protein 6<br>SNF8 – ESCRT-II complex subunit SNF8<br>VPS28 – Vacuolar protein sorting 28<br>VPS35 – Vacuolar protein sorting 35<br>VPS4A – Vacuolar protein sorting 4 homolog A<br>BROX – BRO1 domain and CAAX motif containing<br>HGS – Hepatocyte growth factor-regulated tyrosine kinase substrate<br>STAM2 – Signal transducing adaptor molecule 2 |
| **Flotillins**<br>FLOT1 – Flotillin 1<br>FLOT2 – Flotillin 2 |
| **Annexins (EV-associated)**<br>ANXA1 – Annexin A1<br>ANXA2 – Annexin A2<br>ANXA3 – Annexin A3<br>ANXA4 – Annexin A4<br>ANXA5 – Annexin A5<br>ANXA6 – Annexin A6<br>ANXA7 – Annexin A7<br>ANXA8 – Annexin A8<br>ANXA11 – Annexin A11 |
| **Heat-shock proteins (enriched in exosomes)**<br>HSPA1A / HSPA1B (HSP70) – Heat shock protein family A (Hsp70) member 1A / 1B<br>HSP90AA1 – Heat shock protein 90 alpha family class A member 1<br>HSP90AB1 – Heat shock protein 90 alpha family class B member 1<br>HSPA9 – Heat shock protein family A (Hsp70) member 9<br>HYOU1 – Hypoxia up-regulated 1 |
| **Membrane / endosome / lysosome proteins**<br>LAMP1 – Lysosomal associated membrane protein 1<br>LAMP2 – Lysosomal associated membrane protein 2<br>CAV1 – Caveolin 1<br>STOM – Stomatin |
| **Rab GTPases (EV trafficking-associated)**<br>RAB5A – RAB5A, member RAS oncogene family<br>RAB5B – RAB5B, member RAS oncogene family<br>RAB5C – RAB5C, member RAS oncogene family<br>RAB7A – RAB7A, member RAS oncogene family<br>RAB8A – RAB8A, member RAS oncogene family<br>RAB8B – RAB8B, member RAS oncogene family<br>RAB10 – RAB10, member RAS oncogene family<br>RAB11A – RAB11A, member RAS oncogene family<br>RAB13 – RAB13, member RAS oncogene family<br>RAB18 – RAB18, member RAS oncogene family<br>RAB21 – RAB21, member RAS oncogene family |

**Table 2.**
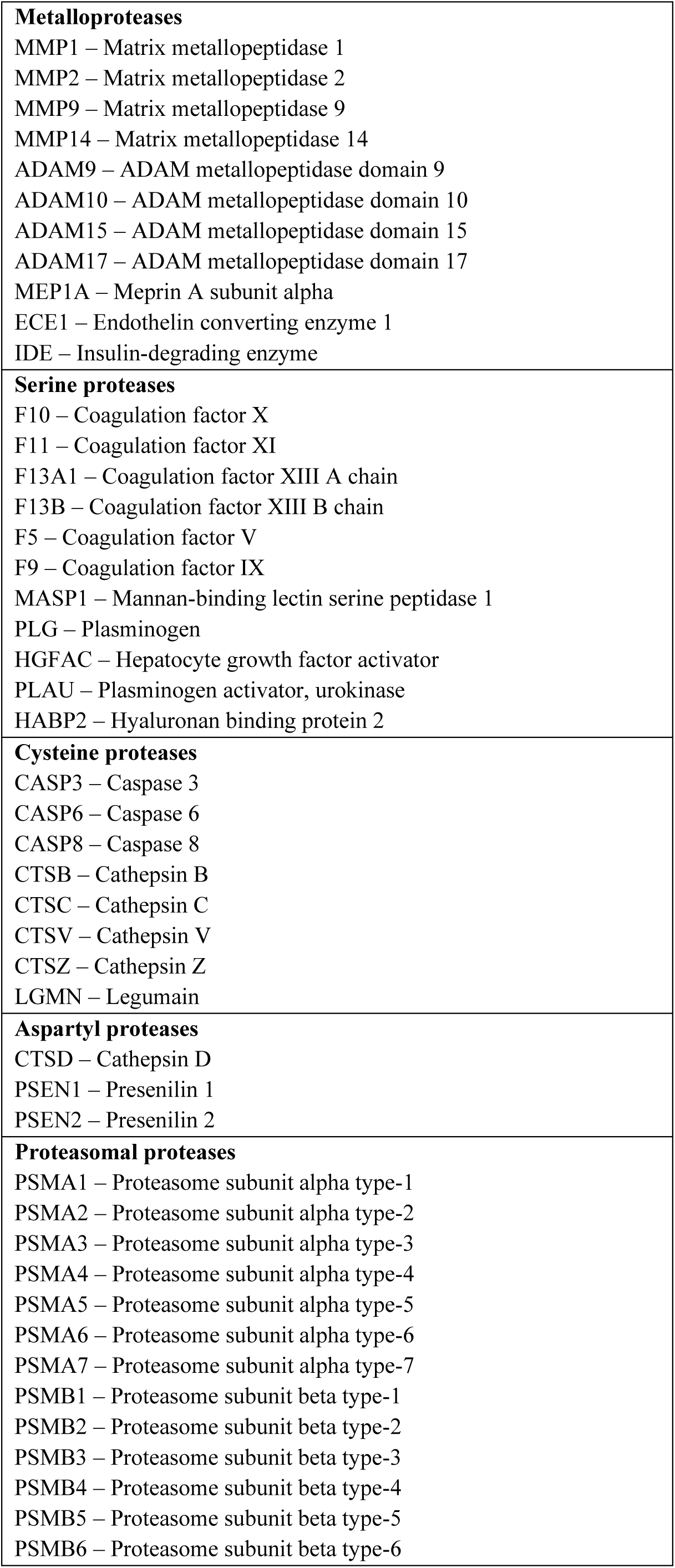
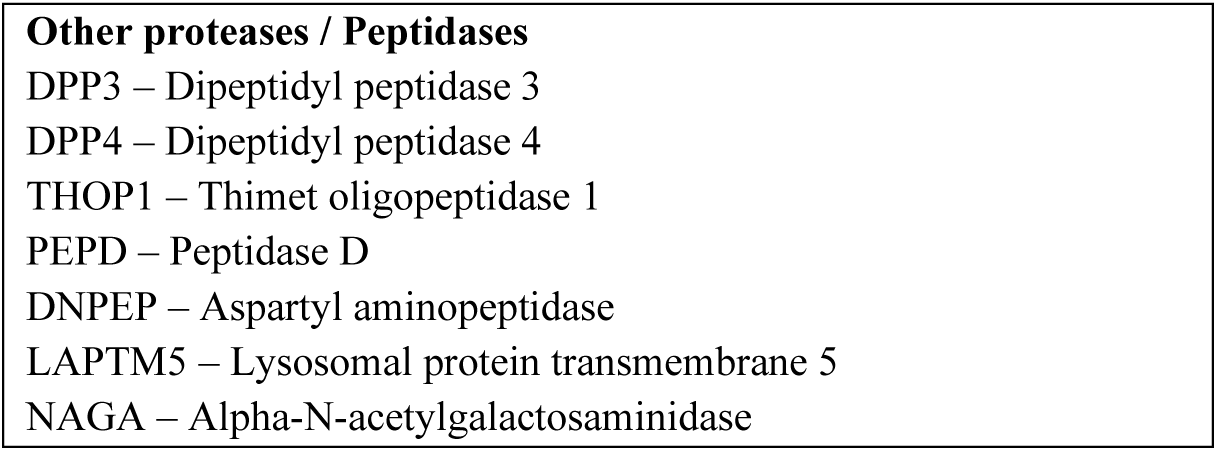
Proteases detected via proteomic analysis of GEC-derived small EVs.

**Table 3.**
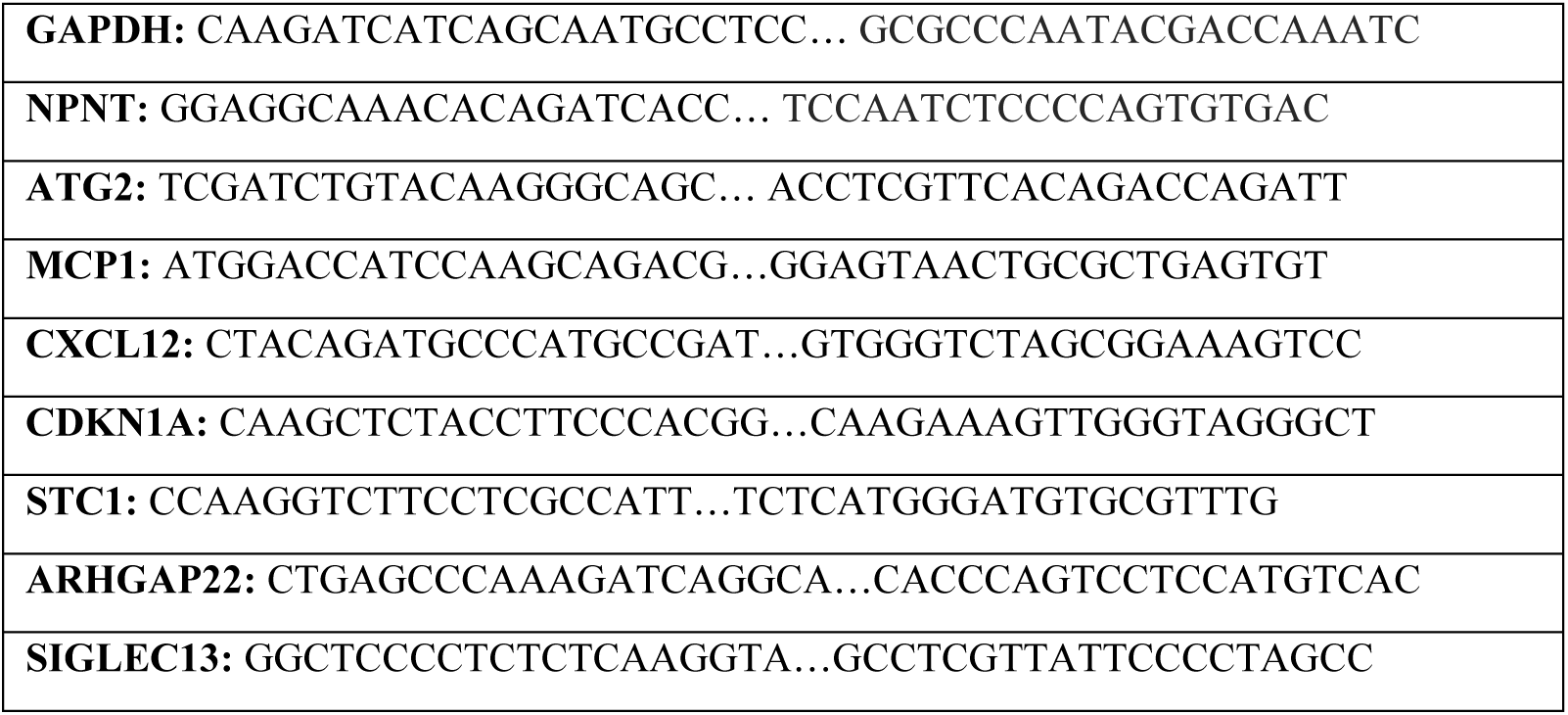
qPCR Primers.

### GEC-derived small EVs display surface protease activity enabling ECM traversal

*In vivo*, GECs and podocytes are separated by the GBM, a dense extracellular barrier. EV-mediated GEC to podocyte signaling therefore requires vesicle passage across this barrier. Proteomic analysis of GEC-derived small EVs revealed the presence of integrins as well as several serine proteases, suggesting that these proteins may facilitate EV interaction with the GBM and support their traversal toward podocytes. To assess whether these surface proteins and proteases are functionally active, we measured proteolytic activity on intact EVs using casein-based assays under surface-preserving conditions. Casein-based protease assays revealed measurable proteolytic activity on GEC-derived EVs which remained constant over time (**Fig. 6A, B**). As collagen is among the most prevalent proteins within the GBM, we assessed the ability of EVs to degrade collagen using a fluorogenic collagenase assay. This method detected collagen cleavage by measuring the increase in fluorescence emitted from a quenched collagen substrate upon proteolytic digestion of GEC-derived small EVs (**Fig. 6C, D**). Using fluorescence resonance energy transfer (FRET)-based probes, we further identified the presence of cathepsins, ADAM10, ADAM13, Calpain, and matrix metalloproteases MMP2 and MMP7 on the EV surface (**Fig. 6E, F**). The activities of these proteases were only partially abolished after heat inactivation. To determine whether this enzymatic activity facilitates ECM penetration, we fluorescently labeled GEC-derived small EVs and incubated them on Matrigel or embedded them in Matrigel, a laminin-, collagen IV-, entactin-, and proteoglycan-rich ECM surrogate mimicking the GBM (**Fig. 6G-J**). Live tracking demonstrated that small EVs migrated in multiple directions and over variable distances within 3 hours when incubated on or in gel, suggesting active movement beyond passive sedimentation (**Fig. 6G-J**). Heat exposure of small EVs inhibited small EV traversal through the matrix (**Fig. 6H, J**). Given that Matrigel pore sizes (∼140 nm) (45) are slightly larger than the median small EV diameter (∼100 nm), we next tested whether small EVs could facilitate the movement of larger complexes. To this end, GEC-derived small EVs were coupled to 4 μm aldehyde-sulfate latex beads via anti-CD63 antibodies, generating complexes that exceed Matrigel pore size. Remarkably, EV-bead complexes also exhibited directional movement within Matrigel, albeit at reduced distance, compared to free EVs (**supplementary Fig. 6**). These findings suggest that protease activity on the small EV surface contributes to ECM remodeling and enables small EV mobility through dense biological barriers.

**Figure 6:**
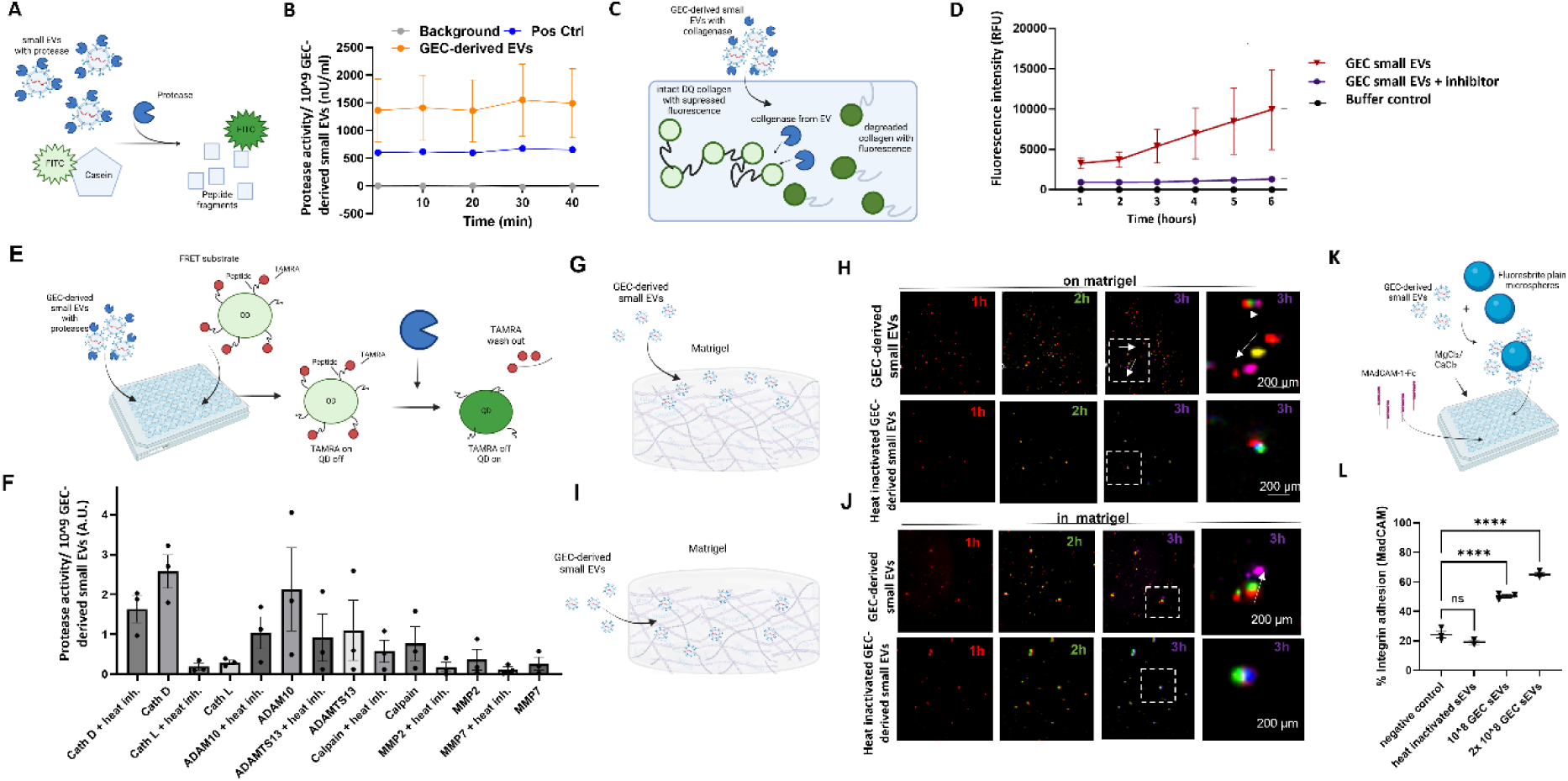
GEC-derived small EVs carry proteases and integrins on their surface that allow them to travel through ECM. A: Schematic illustration of Casein-based proteases activity assay to measure proteases activity of GEC-derived small EVs. B: Quantification of proteases activity of GEC-derived small EVs. Activity is given as total proteases activity per 10^9^ small EVs in nU/mL. Data is given as mean +/- SEM; n = 4. C: Schematic illustration of the in-gel collagenase assay. Small EVs isolated from GECs were incubated with a fluorescently quenched collagen substrate embedded within a polyacrylamide gel matrix (DQ-collagen). Upon proteolytic cleavage by small EV-associated collagenases and other matrix-degrading enzymes, the quenched fluorophore is released and becomes fluorescent, generating a spatially confined signal within the gel. Fluorescence intensity was monitored over time to quantify collagen degradation kinetics. D: Fluorescent intensity resulting from collagenase activity of GEC-derived small EVs measured in the in-gel collagen degradation assay over time. Fluorescent signal reflects progressive cleavage of the collagen-embedded substrate, with increasing intensity corresponding to higher enzymatic activity. Small EV-treated gels were compared to small EV treated with protease inhibitor and buffer-only controls to distinguish active degradation from background fluorescence. * p< 0.05, **** p< 0.0001. E: Schematic illustration of the fluorescence resonance energy transfer (FRET)-based assay used to detect proteolytic activity of cathepsin B, cathepsin D, cathepsin L, ADAM19, ADAM13, calpain, MMP2, and MMP7 on the surface of GEC-derived small EVs. Protease-specific FRET substrates emit fluorescence upon cleavage, enabling detection of active enzymes under surface-preserving conditions. F: Quantification of protease activities measured by the FRET assay shown in panel I. Data are expressed as arbitrary units (AU) normalized to 10⁹ small EVs and presented as mean ± SEM; n = 3. Each measurement was performed with regularly isolated GEC-derived small EVs and small EVs exposed to heat. G: Schematic illustration of small EVs incubated on Matrigel. Fluorescently labeled GEC-derived small EVs were incubated on Matrigel, a laminin- and collagen IV-rich extracellular matrix surrogate that mimics key biochemical and biophysical features of the glomerular basement membrane (GBM). H: Tracking of fluorescently labeled GEC-derived small EVs, with and heat inactivation, incubated on Matrigel over time demonstrates their surface protease–dependent mobility. Red fluorescently labeled small EVs were applied to Matrigel and imaged every hour. For visualization of displacement, small EV signals from the 2 h time point were pseudocolored in green and those from the 3 h time point in magenta, enabling clear distinction of temporal positions in overlay images. The rightmost column shows a magnified view of the 3 h image, with arrows indicating the direction of small EV movement. Scale bar: 200 µm. I: Schematic illustration of GEC-derived small EVs embedded within the Matrigel matrix, depicting their interaction with the dense extracellular network and the concept of protease-supported movement through ECM. J: Tracking of fluorescently labeled GEC-derived small EVs embedded within Matrigel with and heat inactivation, confirming the contribution of small EV-associated protease activity to ECM traversal. As in panel D, red EVs were imaged hourly; signals from 2 h and 3 h were pseudocolored in green and magenta, respectively, to visualize temporal displacement. The rightmost column provides a magnified view of the 3 h image with arrows marking individual small EV trajectories. Scale bar: 200 µm. K: Schematic illustration of the integrin binding assay. Approximately 10¹⁰ GEC-derived small EVs, 2×10¹⁰ GEC-derived small EVs or ¹⁰ GEC-derived heat inactivated small EVs were adsorbed onto 3 µm Fluoresbrite microspheres and incubated in MAdCAM-1-coated 96-well V-bottom plates in the presence of Mg²⁺/Ca²⁺ (integrin-permissive conditions) or EDTA (background control). After 20 min, non-adherent microspheres were quantified using a plate reader, and the percentage of bound microspheres was calculated. L: Quantification of the GEC-derived small EV adhesion assay described in panel K. Data are shown as percentage of bound microspheres; n = 3. Abbreviations: ADAM: a disintegrin and metalloproteinase, ECM: extracellular matrix, EVs: extracellular vesicles, FRET: fluorescence resonance energy transfer, GEC: glomerular endothelial cells, h: hour, miR: microRNA, SEM: standard error of mean.

Integrin-mediated binding likely cooperates with protease activity to facilitate small EV-GBM interactions. Integrin-binding assays confirmed the dose-dependent ability of GEC-derived small EVs to adhere to surfaces which was lost after exposure of GEC-derived small EVs to heat (**Fig. 6K, L**), suggesting that integrins anchor small EVs to extracellular substrates and provide localized contact points that may enhance protease-mediated ECM remodeling and directional traversal. Together, these results indicate that GEC-derived small EVs employ surface protease activity coupled with integrin-mediated binding to overcome the physical barrier of the GBM and enable targeted delivery to podocytes.

### Small EVs traverse the glomerular basement membrane and reach podocytes *in vivo*

To determine whether circulating small EVs can cross the GFB *in vivo*, red fluorescently labelled small EVs derived from cell culture were injected into the circulation of Tg(wt1b:eGFP) zebrafish larvae at 3 days post fertilization (dpf) in control condition (1 % DMSO treatment or CTRL-MO, after PAN induced glomerular damage or after morpholino-mediated *cd2ap* knockdown (**Fig. 7A**). In this transgenic line, GFP expression labels podocytes and renal tubular epithelial cells, whereas GECs remain unlabeled, enabling spatial resolution of glomerular compartments. A schematic overview of the anatomy of the pronephric glomerulus which fuses at the midline and drains into paired tubular segments is shown in **Fig. 7B**. Confocal microscopy of the pronephric region with three-dimensional reconstruction revealed accumulation of small EVs within the glomerulus, with no detectable signal in the tubular compartment, indicating preserved filtration barrier integrity under baseline conditions (**supplementary Fig. 7**). Consistent with these findings, confocal imaging of fixed pronephroi isolated 24 hours after small EV injection demonstrated red fluorescent small EVs localized within the podocyte region (**Fig. 7C**, white arrows), while remaining absent from tubular epithelial cells and the tubular lumen. In contrast, following PAN-induced glomerular injury, small EVs were detected not only in podocytes but also within proximal tubular epithelial cells (**Fig. 7D**), suggesting compromised barrier function. Cryosection analysis provided further evidence for transglomerular passage of small EVs. Notably, red fluorescent small EVs colocalized with GFP-positive podocytes, but were not observed in proximal tubular epithelial cells under baseline conditions (**Fig. 7E**). This selective localization indicated successful traversal of the GBM with preferential delivery to podocytes. However, following podocyte injury induced by *cd2ap* morpholino injection at the one-cell stage, small EVs were additionally detected in proximal tubular cells (**Fig. 7F**).

**Figure 7:**
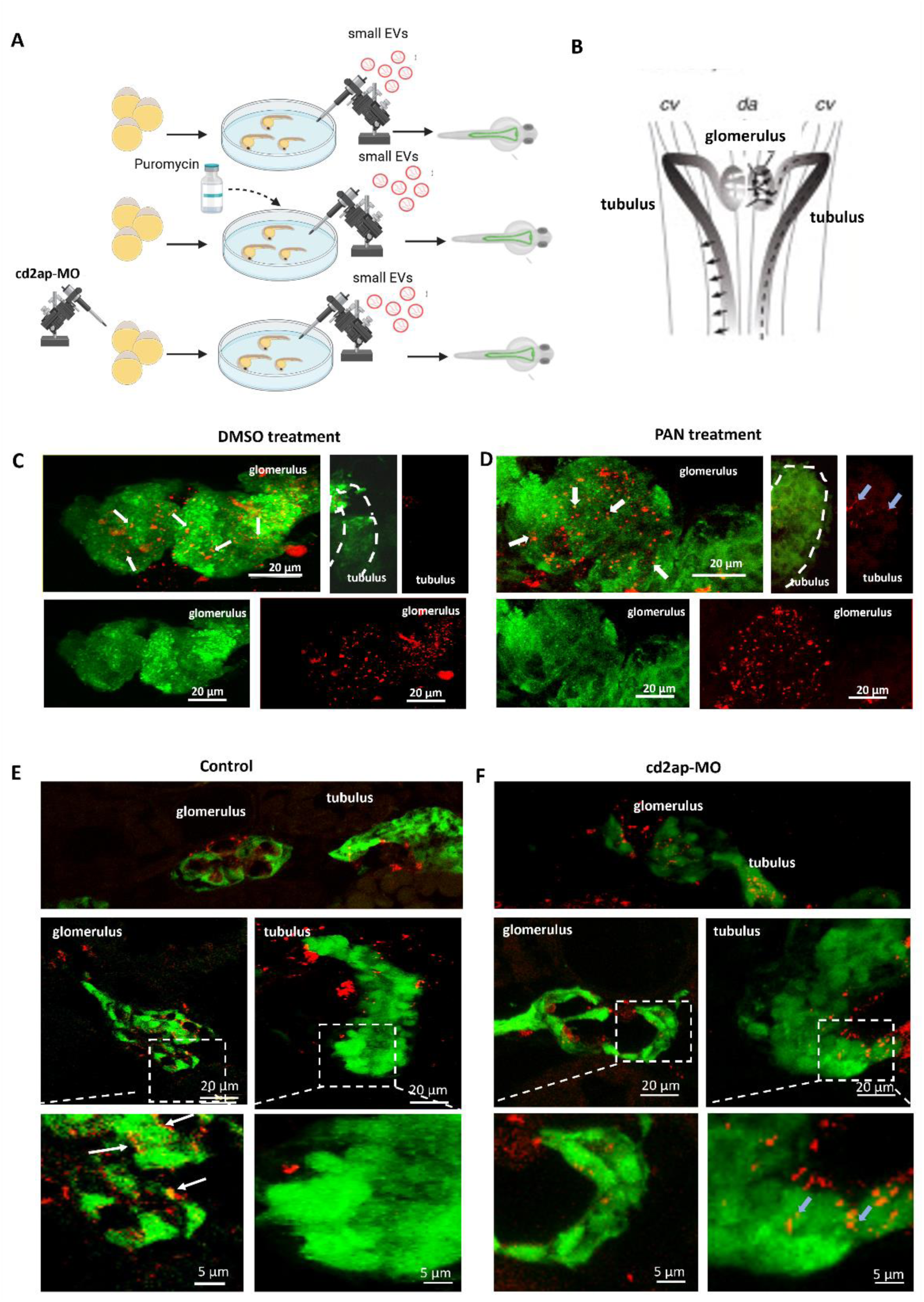
*S*mall EVs can pass the glomerular basement membrane in zebrafish. A: Experimental setup: Small EVs from human cell culture supernatant were labeled with a red dye (PHK 26) and injected into Tg(wt1b:eGFP) zebrafish larvae at 72 hpf. Zebrafish were treated with DMSO or treated with puromycin (PAN) at 46 hpf or injected with cd2ap morpholino or CTRL-MO at egg stage. B: Schematic of the zebrafish pronephros at 4 dpf, showing a fused glomerulus and two tubular segments. In Tg(wt1b:eGFP) larvae, podocytes and tubule cells are green; glomerular endothelial cells are unlabeled. C/D: Confocal images of pronephroi isolated from DMSO control treated zebrafish (C) and from zebrafish treated with puromycin (PAN) (D) that were injected with red fluorescent small EVs in the circulation at 72 hpf. Red fluorescent small EVs were overlapping with green fluorescent signal of podocytes (white arrows). Small EVs were not detectable in proximal tubular cells in control zebrafish but overlapped with tubular signal in PAN treated zebrafish (blue arrows). Scale bar 20 µm. Note that some small EVs formed aggregates. E/F: Cryosections of control (CTRL-MO injected, first row and untreated in the other pictures) Tg(wt1b:eGFP) zebrafish (F) and of Tg(wt1b:eGFP) zebrafish with morpholino induced *cd2ap* knockdown (F) that were injected with red fluorescent small EVs in the circulation at 72 hpf and imaged 24 hours later. Scale bar 20 µm. Tubular and glomerular region are zoomed in in the right picture. Red fluorescent small EVs were overlapping with green fluorescent signal of podocytes (white arrows). Small EVs were not detectable in proximal tubular cells in control zebrafish (CTRL-MO) but overlapped with tubular signal in *cd2ap* knockdown zebrafish (blue arrows). Scale bar 20 µm and 5 µm. Note that some small EVs formed aggregates. Abbreviations: CTRL: control, dpf: day post fertilization, EVs: extracellular vesicles, h: hours, hpf: hours post fertilization, MO: morpholino, PAN: puromycin

Together, these data suggest that circulating small EVs can cross the GBM in zebrafish larvae and selectively interact with podocytes under physiological conditions, while gaining access to the tubular compartment is only possible upon glomerular injury.

## Discussion

Small EVs are increasingly recognized as key mediators of intercellular communication, contributing to both physiological homeostasis and disease progression. In the kidney, small EVs are secreted by a variety of resident cell types, including podocytes, tubular epithelial cells, mesangial cells, and GECs. Small EVs are known to carry diverse cargo such as proteins, lipids and nucleic acids, which can influence target cell behavior. These vesicles have been implicated in processes ranging from immune modulation and fibrosis to GFB integrity and tubular cross-talk. However, most mechanistic insights, such as molecular composition, biogenesis and functional relevance, have largely relied on *in vitro* systems (46, 47), leaving open critical questions regarding their physiological role and complexity in the native renal microenvironment.

A long-standing question in the field of glomerular biology is the feasibility of communication between GECs and podocytes, two cell types which are spatially separated by the size selective GBM. The GBM serves both as a structural and functional unit in the glomerular filtration barrier with pores estimated to be in the range of 2-8 nm and presumed to restrict the passage of small EVs (48, 49). It has therefore been unclear whether GEC-to-podocyte communication via small EVs occurs under such biophysiologically constrained conditions. However, using an integrated approach combining GEC-derived small EV-omics, podocyte transcriptomics, and *in vivo* imaging in zebrafish, we demonstrate that GEC-derived small EVs can cross the GBM and functionally modulate podocyte gene expression.

We first established that GECs secrete small EVs consistent with exosome characteristics, including a median diameter of 100-150 nm, canonical morphology, and enrichment of tetraspanins CD63, CD9, and CD81 and absence of Calcein. The incorporation of SRS microscopy in our study provided a label-free, chemically specific approach to verify the molecular integrity of GEC-derived small EVs. SRS enabled interrogation of the CH-stretching region (≈2850–2950 cm⁻¹) and the fingerprint region (≈600-1800 cm⁻¹) that are two key vibrational domains for EV characterization (50). The spectral features of GEC-derived small EVs were consistent with previously reported Raman-based characterizations of EVs (51). The dominance of CH₂ over CH₃ vibrations was characteristic of lipid bilayer membranes and were consistent with the molecular composition of exosomes (52). Overall, these SRS measurements confirmed the presence of vesicle-associated lipid and protein signatures, with the CH-stretch region providing the most robust and reliable contrast for exosome detection under the current experimental conditions. The ability of SRS to simultaneously resolve these lipid- and protein-associated vibrational signatures underscores its strength as a non-destructive, label-free tool for EV profiling. Moreover, correlative fluorescence-SRS imaging ensured that the Raman-active structures corresponded to bona fide GEC-derived small EVs, thereby enhancing confidence in their biochemical annotation. These findings demonstrate that SRS microscopy offers unique molecular-level insights into small EV composition, complementing conventional biochemical assays and strengthening our interpretation of GEC small EV identity and function within the glomerular microenvironment. Endosomal origin of small EVs was confirmed using a pH-sensitive reporter construct, showing selective secretion of CD63-positive vesicles (53).

RNA sequencing revealed that miRs represented the major fraction of mapped reads of GEC-derived small EV cargo. Among these, miR-192, miR-143, and miR-26b were selectively enriched in GEC-derived small EVs. These miR families have been shown to play important roles in GEC to podocyte communication before (15, 54–56) and our data is the first to show that they are packed into small EVs. By combining absolute quantification of small EV miR cargo with functional transcriptomic profiling of recipient podocytes, we demonstrate that GEC-derived small EVs deliver selective miRs which impact on podocyte transcriptional networks. Upregulation of genes such as *STC1* and *ARHGAP22*, together with downregulation of inflammatory mediators like *MCP1*, points to a shift away from maladaptive inflammatory signaling toward cytoskeletal stabilization and adaptive stress responses in podocytes due to GEC-derived small EVs. Integrative analyses revealed that many of these transcriptional effects could be attributed to direct actions of transferred small EV miRs. Notably, miR-192 of GEC-derived small EV was predicted and experimentally validated to target *NPNT*, an extracellular matrix protein essential for GBM integrity, and *ATG2* an autophagy regulator in podocyte. Importantly, these interactions were not restricted to artificial settings of isolated vesicle transfer. Using a co-culture transwell model and direct visualization of fluorescently labeled miRs within EVs, we demonstrated GEC to podocyte transfer of miR cargo under conditions mimicking physiological exchange. This is in line with previously reported data, that small EVs act as bona fide messengers complementing classical paracrine signaling mechanisms (53, 57). Taken together, our findings hint GEC-derived small EVs as a central regulatory axis in glomerular homeostasis. By delivering a tailored miR cargo, GECs modulate podocyte transcriptional programs which converge on inflammation, cytoskeletal organization, extracellular matrix remodeling, and autophagy.

However *in vivo*, GECs and podocytes are separated by the GBM raising the question whether small EV mediated communication from GECs to podocytes could actually happen *in vivo*. The GBM is a dense extracellular matrix composed of laminins, collagen IV, nidogens, and heparan sulfate proteoglycans (58). Proteolytic enzymes have previously been implicated in GBM remodeling and podocyte injury (59). Proteomic profiling of GEC-derived small EVs revealed not only a repertoire enriched in classical exosome markers but also proteases and integrins, indicating that these vesicles are equipped to remodel the GBM, interact with podocytes, and potentially facilitate barrier crossing via extracellular matrix engagement. In independent experiments we showed that GEC-derived small EVs carry active surface proteases, including serine proteases, cathepsins, ADAM10, ADAM13, Calpain, and matrix metalloproteases (MMP2 and MMP7), as well as integrins, which together enable small EV-matrix interactions and remodeling. Fluorescence-based tracking in Matrigel demonstrated that small EVs can migrate actively within ECM-like environments, and even mediate the movement of larger complexes exceeding the Matrigel pore size. These findings highlight a dual strategy by which small EVs employ both proteolytic activity and integrin-mediated adhesion to overcome dense biological barriers. This mechanism provides a plausible explanation for how small EVs overcome the GBM barrier to deliver functional cargo to podocytes, extending previous observations from tumor and fibrotic models (60). Our protease activity assays revealed that heat inactivation only partially reduces enzymatic activity. This observation is consistent with several key factors. First, exosomes can protect their internal cargo enzymes, allowing internal proteases to remain active despite heat treatment (61). Second, some proteases exhibit unusual heat stability or can regain partial structure upon cooling. For example, metalloproteases can maintain partial structural integrity if the temperature is not sufficiently high, serine proteases, though generally heat-labile, can retain low residual activity, and cysteine proteases vary in stability, with some capacity of partial refolding after moderate heating (61). Finally, small EV aggregation under heat treatment may trap proteases within partially protected microdomains, further contributing to residual activity (62). Integrins on the small EV surface likely cooperate with protease activity to facilitate localized ECM remodeling. By anchoring small EVs to ECM substrates, integrins may enhance the efficiency of directional traversal, providing spatially confined sites where proteases can act on the GBM. This mechanism may be particularly important in the glomerular context, where the GBM represents a highly dense and selective barrier (63). Our findings therefore suggest that small EV-mediated intercellular communication is not a passive process but a coordinated interplay between surface adhesion molecules and enzymatic activity. The ability of small EVs to transport enzymes capable of remodeling dense ECM has broader implications for tissue-specific communication and targeted cargo delivery. These findings expand the current paradigm of intercellular communication within the glomerulus and have implications for the design of small EV-based therapeutics targeting renal or other dense tissue barriers.

Previous work has established the role of podocyte-derived vesicles in paracrine signaling and injury propagation (64, 65). However, whether circulating vesicles can traverse the GFB and interact with podocytes *in vivo* has not been conclusively demonstrated. Using an *in vivo* zebrafish model with cell type-resolved labeling of podocytes and proximal tubular cells, we demonstrate that circulating small EVs are capable of traversing the GBM and selectively localize to podocytes in the intact glomerulus. The absence of small EVs in tubular cells under baseline conditions suggests that the GFB was intact and that vesicle passage is not a mere artifact of barrier breakdown. In contrast, disruption of the filtration barrier either by PAN-induced injury or by perturbation of podocyte slit diaphragm integrity through *cd2AP* knockdown permitted small EV access to the tubular compartment. Small EV transversal across the GBM in healthy as well as diseased conditions was most likely facilitated by protease activity on small EVs. This finding challenges the traditional view of the GBM as an impermeable barrier to nanoscale particles and supports the concept of bidirectional vesicle-mediated communication within the glomerular filtration unit.

Taken together, these findings point toward a model where protease-mediated micro-remodeling of the extracellular matrix by small EVs allows their passage through the GBM, followed by integrin-dependent docking and uptake by podocytes. This model highlights small EVs not only as carriers of signaling molecules but also as active biological agents capable of shaping their microenvironment to deliver cargo across restrictive barriers. Future studies in mammalian models will be critical to determine whether similar protease- and integrin-mediated mechanisms operate in human kidney disease.

## Conclusion

This study provides first *in vivo* evidence that GEC-derived small EVs can traverse the GBM and deliver functional cargo to podocytes. We identify vesicle-associated proteases as facilitators of GBM passage and integrin-dependent interactions as potential determinants of podocyte targeting. The selective miR and protein cargo of these vesicles suggests a mechanism by which small EVs modulate podocyte transcriptional and extracellular matrix programs under both physiological and stress conditions. In glomerular disease, systemic and GEC-derived small EVs can transverse not only the GBM but also the GFB, delivering cargo to tubular compartments. Collectively, our findings redefine the GBM from a passive filter to a site of active vesicle-mediated communication and identify GEC-derived small EVs as potential therapeutic targets in kidney disease.

## Supporting information

Supplementary report

## Author Contributions

ST: Conceptualization, Data curation, Formal analysis, Methodology, Software, Writing – review and editing; JK: Conceptualization, Data curation, Formal analysis, Methodology, Software, Writing – review and editing; NS: Methodology, Data curation, Validation, Formal analysis, Writing – review and editing; AO: Methodology, Validation; PL: Data curation, Formal analysis, Writing – review and editing; AB: Resources, Writing – review and editing; JVD: Data curation, Formal analysis; MK: Data curation, Formal analysis, Writing – review and editing; GS: Data curation, Formal analysis; SC: Resources; CD: Methodology, Writing – review and editing; MS: Resource, Writing – review and editing; SU: Conceptualization, Data curation, Funding acquisition, Formal analysis, Writing – review and editing; JMD: Conceptualization, Data curation, Funding acquisition, Investigation, Project administration, Resources, Supervision, Visualization, Writing – original draft, Writing – review and editing.

## Data Sharing Statement

The mass spectrometry proteomics data have been deposited to the ProteomeXchange Consortium via the PRIDE (66) partner repository with the dataset identifier PXD073057.

