## Supplementray material for "Glomerular Endothelial Cell-Derived Extracellular Vesicles Cross the Basement Membrane to Regulate Podocyte Function"

### Supplemental Material

#### Table of content:

Supplementary Figure 1

Supplementary Figure 2

Supplementary Figure 3

Supplementary Figure 4

Supplementary Figure 5

Supplementary Figure 6

Supplementary Figure 7

Supplementary table 1

Supplementary table 2

Supplementary table 3

Supplementary data

Supplementary video 1

#### Supplementary Figure legend

**Supplementary Figure 1:** ***Composition of RNA detected in GEC- and podocyte-derived small EVs***

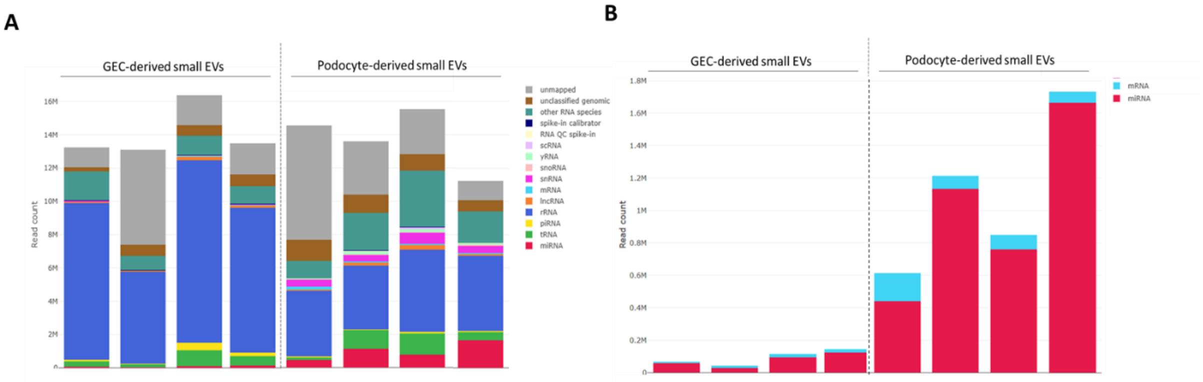

A: Relative read composition of RNA detected in GEC- and podocyte-derived small EVs. After processing of the reads (adapter trimming, quality filtering, size filtering), all remaining reads were mapped against various databases to categorize them. This is done in a hierarchical process, where reads are first mapped against the genome. “Unclassified genomic” indicates reads that were mapped against the genome but were not found in any of the RNA specific databases, while “unmapped” are reads that could not be found in the given reference genome.

B: Absolut read counts of mRNAs and miRs only detected in GEC- and podocyte-derived small EVs.

Abbreviations: EVs: extracellular vesicles, GEC: glomerular endothelial cells, miR: microRNA.

**Supplementary Figure 2:** ***miR profile of GEC- and podocyte-derived small EVs.***

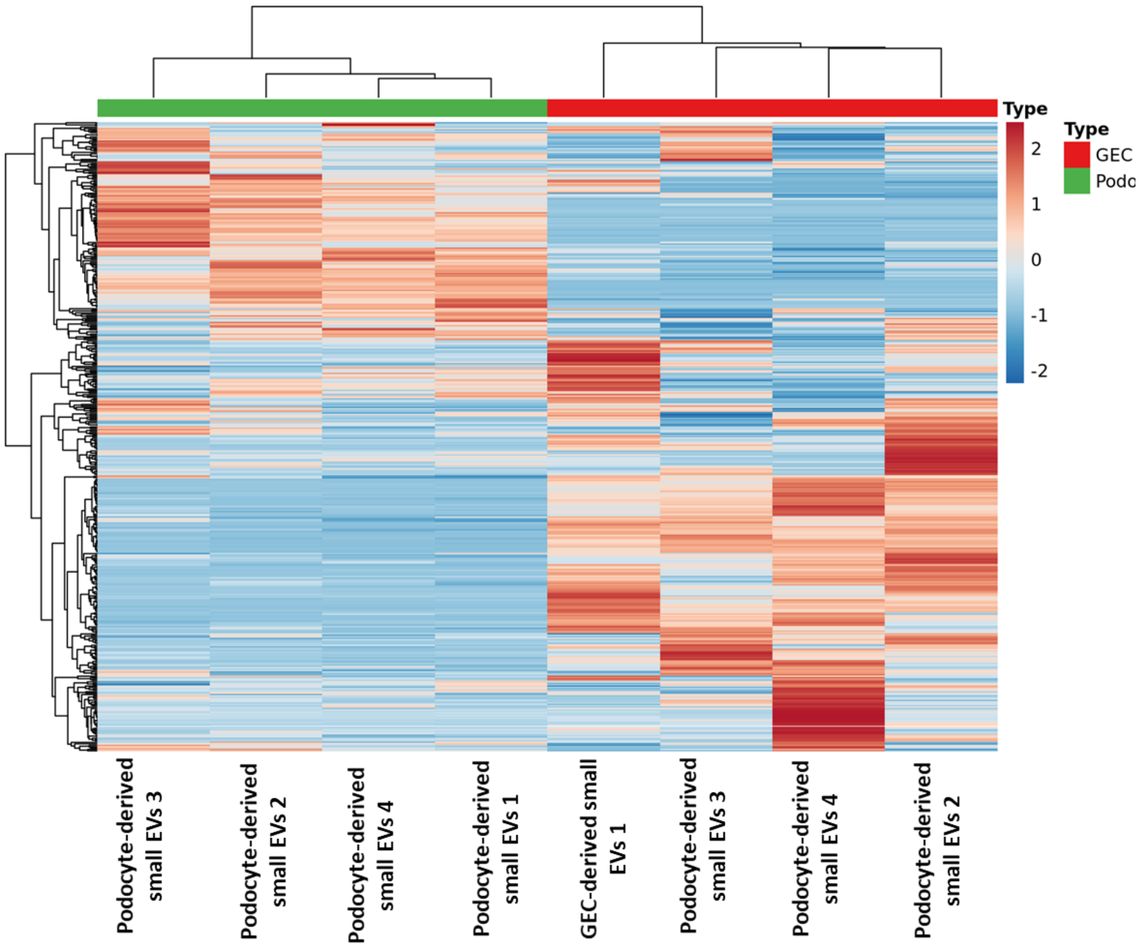


Heatmap visualizes the 447 miRs detectable in GEC and podocyte-derived small EVs.
Data is based on RPM normalized reads and scaled using the unit variance method for visualization in heatmaps. Clustering was done using the average method of heatmap calculating the distances as correlations. To avoid spurious candidates the following presence threshold was applied: each miR must have had RPM > 5 in at least 1/n (groups) of the samples. This ensures to capture miRs with both variability and sufficient biological representation.

columns = GEC and podocyte small EVs, rows = miRs, color scale = relative expression (blue = low, red = high), top dendrogram = sample clustering, left dendrogram = hierarchical clustering of miRs.

**Supplementary Figure 3:** ***Podocyte mRNA regulated after treatment with GEC-derived small EVs.***

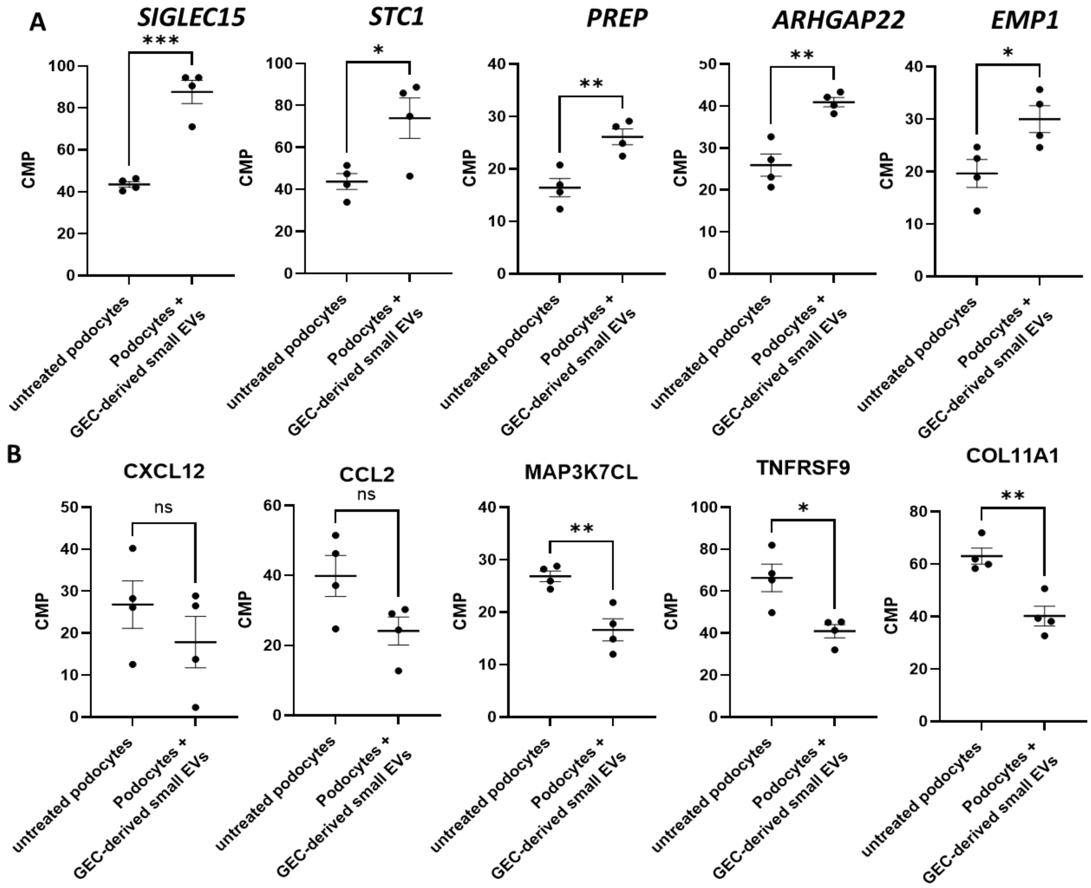


A: Top 5 upregulated mRNAs in podocytes treated with GEC-derived small EVs compared to untreated podocytes (FDR < 0.05 only). mRNAs are given with their CPM values. CPM normalized values are calculated by edgeR. n = 4. single CPM values, mean and SEM are given. * p< 0.05, ** p<0.01, *** p<0,001.

B: Top 5 downregulated mRNAs in podocytes treated with GEC-derived small EVs compared to untreated podocytes (FDR < 0.05 only) with their CPM values. CPM normalized values are calculated by edgeR. n = 4. single CPM values, mean and SEM are given. n.s.: non-significant, * p< 0.05, ** p<0.01.

Abbreviations: CPM: counts per million, EVs: extracellular vesicles, FDR: False discovery rate, GEC: glomerular endothelial cells, SEM: standard error of mean.

**Supplementary Figure 4:** ***GO enrichment analysis of podocyte mRNA regulated after treatment with GEC-derived small EVs.***

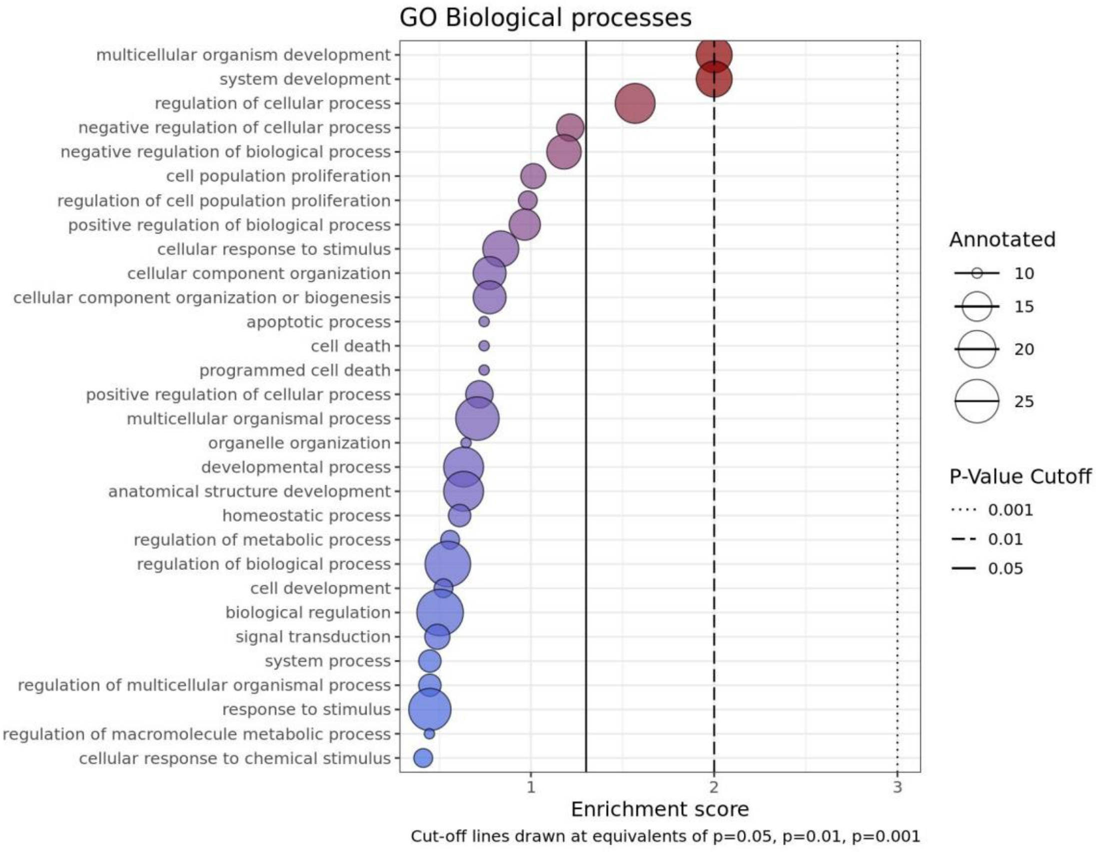


Differentially expressed mRNAs based on a FDR cutoff (FDR <= 0.05) were used in a GO-term enrichment analysis. Enriched biological processes (BP) were identified by using the Kolmogorov Smirnov (KS) test. The 30 enriched BPs are visualized. Dots present the number of genes in the BP as the size of the dots. The enrichment score is shown as -log(KS) on the x-axis as well as indicated by the color code. Lines are drawn at equivalents to p = 0.01, p = 0.05 and p = 0.1.

**Supplementary Figure 5:** ***In silico binding between miR-192 and NPNT as well as miR-192 and ATG2.***

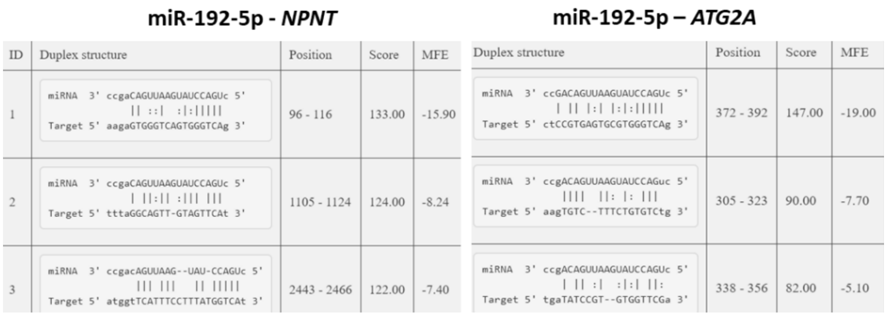

**miR-192-5p - *NPNT***

<table>
<tr><th>ID</th><th>Duplex structure</th><th>Position</th><th>Score</th><th>MFE</th></tr>
<tr><td>1</td><td>miRNA 3' ccgaCAGUUAAGUAUCCAGUc 5'<br>|| ::| :|:|||||<br>Target 5' aagaGTGGGTCAGTGGGTCAg 3'</td><td>96 - 116</td><td>133.00</td><td>-15.90</td></tr>
<tr><td>2</td><td>miRNA 3' ccgaCAGUUAAGUAUCCAGUc 5'<br>| ||:|| :||| |||<br>Target 5' tttaGGCAGTT-GTAGTTCAt 3'</td><td>1105 - 1124</td><td>124.00</td><td>-8.24</td></tr>
<tr><td>3</td><td>miRNA 3' ccgacAGUUAAG--UAU-CCAGUc 5'<br>||| ||| || |||||<br>Target 5' atggtTCATTTCCTTTATGGTCAt 3'</td><td>2443 - 2466</td><td>122.00</td><td>-7.40</td></tr>
</table>

**miR-192-5p – *ATG2A***

<table>
<tr><th>Duplex structure</th><th>Position</th><th>Score</th><th>MFE</th></tr>
<tr><td>miRNA 3' ccGACAGUUAAGUAUCCAGUc 5'<br>| || |:| |:|:|||||<br>Target 5' ctCCGTGAGTGCGTGGGTCAg 3'</td><td>372 - 392</td><td>147.00</td><td>-19.00</td></tr>
<tr><td>miRNA 3' ccgACAGUUAAGUAUCCAGuc 5'<br>|||| ||: |: |||<br>Target 5' aagTGTC--TTTCTGTGTCtg 3'</td><td>305 - 323</td><td>90.00</td><td>-7.70</td></tr>
<tr><td>miRNA 3' ccgACAGUUAAGUAUCCAGUc 5'<br>| || :| :|:| ||:<br>Target 5' tgaTATCCGT--GTGGTTCGa 3'</td><td>338 - 356</td><td>82.00</td><td>-5.10</td></tr>
</table>

*In silico* binding was predicted with miRanda.

**Supplementary Figure 6: Small *EV traveling through matrices.***

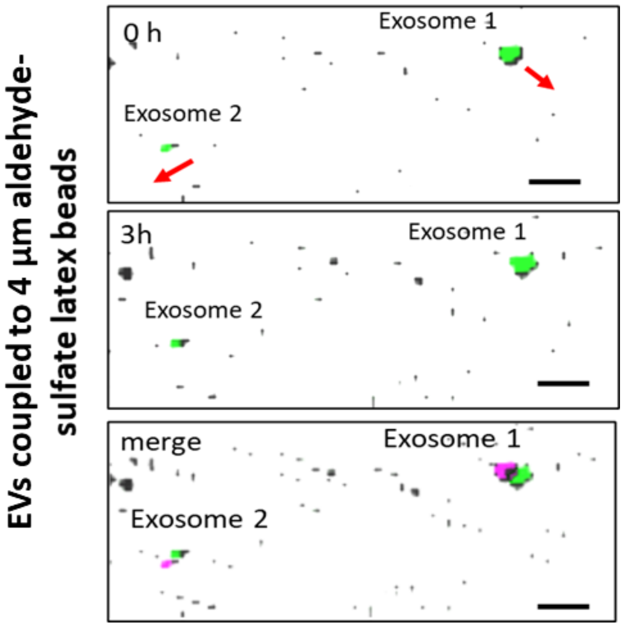


A: Tracking of fluorescently labeled GEC-derived small EVs bound to 4 µm aldehyde-sulfate latex beads in Matrigel over time shows the functionality of their protease activity. The same small EVs are imaged at baseline (0 h) shown with pseudo color green and after 3h shown in pseudo color magenta. Scale bar 10 µm.

Abbreviations: EVs: extracellular vesicles, GEC: glomerular endothelial cells, h: hour.

**Supplementary Figure 7: *Multiphoton microscopy of pronephros of Tg(wt1b:eGFP) zebrafish larvae injected with small EVs.***

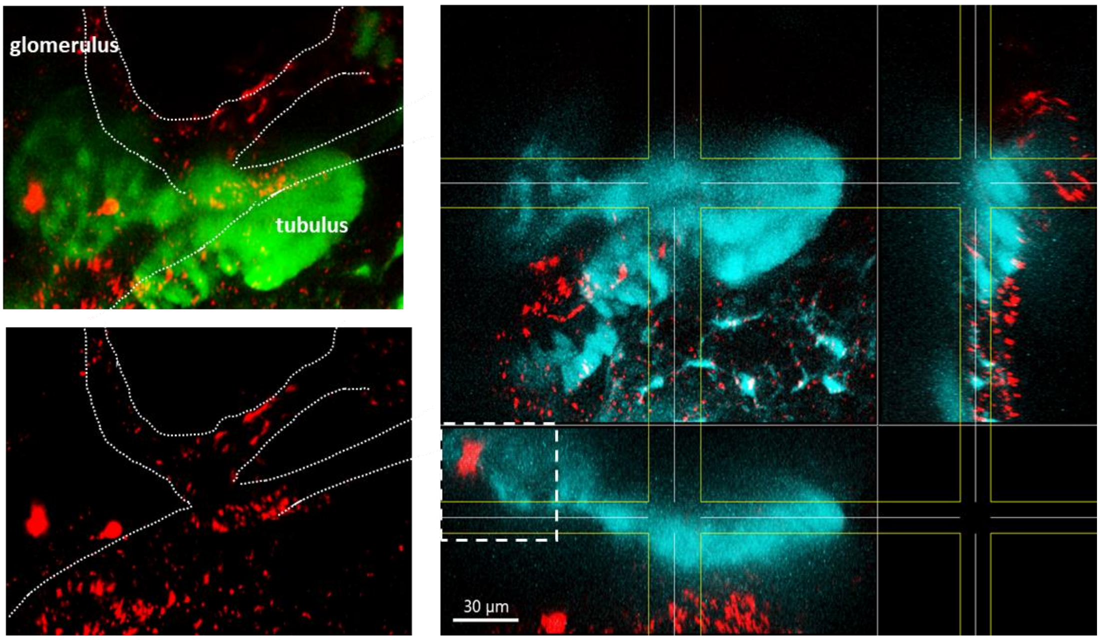

Multiphoton microscopy of pronephros of Tg(wt1b:eGFP) zebrafish larvae at 4 dpf. Red fluorescent small EVs were injected into the zebrafish circulation at 72 hpf. Images were taken 24 h after injection. Reconstruction of the image illustrated that red fluorescent small EVs did not reach tubular cells/ tubular lumen when the glomerular filtration barrier is intact. Small EVs were in vessels next to the tubular system.

Abbreviations: dpf: days post fertilization, hpf: hours post fertilization, EVs: extracellular vesicles.

**Supplementary tables**

**Supplementary table 1**

***Top 100 miRs detectable in GEC-derived small EVs. (Excel file)***

**Supplementary table 2**

***Top 100 miRs detectable in podocyte-derived small EVs. (Excel file)***

**Supplementary table 3**

***Proteins in GEC-derived small EVs reproducibly detectable 5 independent batches of small EVs. (Excel file)***

**Supplementary data**
***multiQC report (multiqc_report.html).***

**Supplementary video 1**
***Z-stack of podocytes in co-culture with GEC that were transfected with CD63 reporter plasmid showing uptake of green fluorescent small EVs in podocytes.***
