## Supplementary figures and images for "Glomerular Endothelial Cell-Derived Extracellular Vesicles Cross the Basement Membrane to Regulate Podocyte Function"

### Supplementary video

#### Slide 1
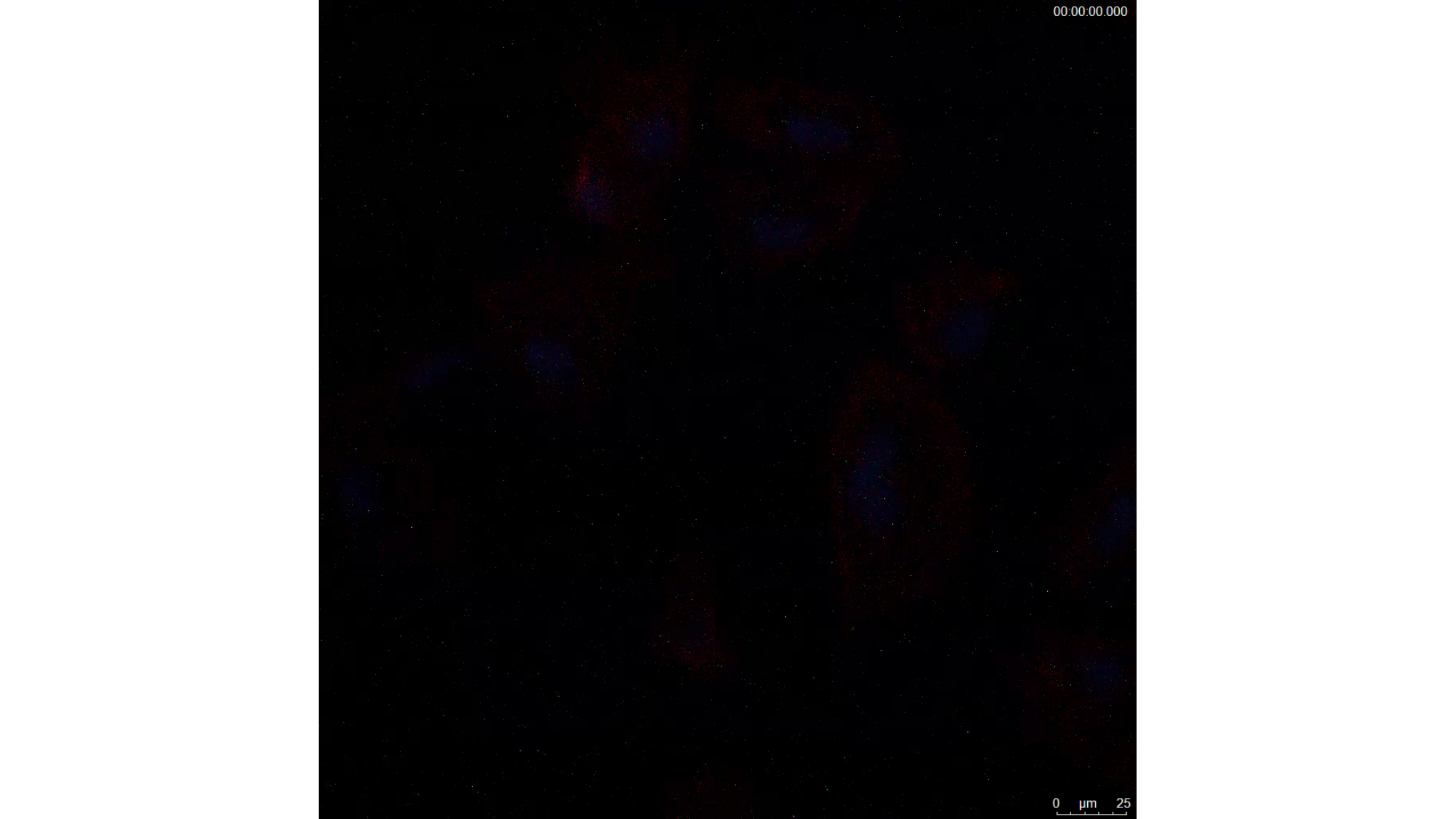
